# Marburg and Ebola virus infections elicit a complex, muted inflammatory state in bats

**DOI:** 10.1101/2020.04.13.039503

**Authors:** Anitha D. Jayaprakash, Adam J. Ronk, Abhishek N. Prasad, Michael F. Covington, Kathryn R. Stein, Toni M. Schwarz, Saboor Hekmaty, Karla A. Fenton, Thomas W. Geisbert, Christopher F. Basler, Alexander Bukreyev, Ravi Sachidanandam

**Author notes:** Email for correspondence: Alexander Bukreyev.

## Abstract

The Marburg and Ebola filoviruses cause a severe, often fatal, disease in humans and nonhuman primates but have only subclinical effects in bats, including Egyptian rousettes, which are a natural reservoir of Marburg virus. A fundamental question is why these viruses are highly pathogenic in humans but fail to cause disease in bats. To understand how bats resist the disease caused by filoviruses, we infected one cohort of Egyptian rousette bats with Marburg virus and another cohort with Ebola virus and harvested multiple tissues for mRNA expression analysis. While virus transcripts were found primarily in the liver, Principal component analysis (PCA) revealed coordinated changes across multiple tissues. Gene signatures in kidney and liver pointed at induction of vasodilation, reduction of coagulation and changes in the regulation of iron metabolism. Signatures of immune response detected in spleen and liver indicated a robust anti-inflammatory state signified by macrophages in the M2 state and an active T cell response. Many of the responsive genes were found to be evolutionarily divergent, providing a framework for understanding the differences in outcomes of filovirus infections between bats and humans. In this study, we outline multiple interconnected pathways that respond to infection by MARV and EBOV, providing insights into the complexity of the mechanisms that enable bats to resist the disease caused by filoviral infections. The results have the potential to aid in the development of new strategies to effectively mitigate and treat the disease caused by these viruses in humans.

## Introduction

The filoviruses Marburg (MARV) and Ebola (EBOV) cause severe and frequently fatal diseases in humans (Rougeron et al., 2015). The diseases caused by these viruses are characterized by hypotension, multisystem organ failure, sepsis-like symptoms, and disseminated intravascular coagulation (DIC) due to profound immune dysregulation accompanied by a cytokine storm (Geisbert et al., 2003). The aggressive use of a recently approved EBOV vaccine has achieved control of the most recent large outbreak in the Democratic Republic of the Congo (Zarocostas, 2020). However, once the disease is manifested, medical intervention can prove to be insufficient, despite the recent progress in the development of antivirals and antibody-based treatments (Dhama et al., 2018; Kortepeter et al., 2020). Therefore, there is an ongoing need to better understand the pathobiology of these viruses. Remarkably, MARV and EBOV appear to cause no significant disease in their confirmed (MARV) or likely (EBOV) bat reservoirs, suggesting that understanding the molecular mechanisms of this resistance can be used to develop therapies for humans.

MARV has been isolated from the Egyptian rousette bat (ERB) (*Rousettus aegyptiacus)* (Towner et al., 2009; Amman et al., 2012, 2020), and ecological and experimental studies have demonstrated that this species serves as a reservoir for the virus (Amman et al., 2012; Schuh et al., 2017a). Experimental infections of ERBs with MARV have consistently demonstrated that despite viral replication in multiple tissues, animals develop a mostly subclinical disease. This is characterized by mild pathology, including transient elevation of alanine aminotransferase, elevated lymphocyte and monocyte counts, and some evidence of minimal inflammatory infiltration in the liver (Paweska et al., 2012, 2015; Amman et al., 2015; Jones et al., 2015, 2019). Transmission has been demonstrated between co-housed ERBs, and the virus is known to be shed in saliva, urine, and feces(Schuh et al., 2017a). However, ERBs do not appear to develop a severe or chronic infection when exposed to MARV, and instead clear the virus and develop at least temporary immunity, including MARV-specific IgG (Schuh et al., 2017b). Circumstantial evidence, including detection of EBOV RNA and anti-EBOV antibodies, suggests that multiple species of bats are also reservoirs for EBOV (Leroy et al., 2009; Marí Saéz et al., 2015). However, infectious EBOV has never been isolated from a wild bat (Dovih et al., 2019). Although serological surveys have identified antibodies reactive to EBOV antigen in ERBs (Pourrut et al., 2009; Yuan et al., 2012; Olival et al., 2013), experimental infection studies performed to date suggest that this bat species is refractory to infection (Krähling et al., 2010), making it an unlikely candidate reservoir for the virus.

The ability of bats to tolerate viral infections has been a topic of considerable interest, and several models have been proposed to explain this phenomenon. One model posits that bats constitutively express interferons to maintain a basal level of innate immune activity (Zhou et al., 2016), although the universality of this model in bats is uncertain (Glennon et al., 2015; Pavlovich et al., 2018). Another model suggests that the resistance of bats to clinical disease is due to a weakened innate immune response, which is attenuated by modifications of some proteins such as the stimulator of interferon genes (STING/TMEM173) (Xie et al., 2018). These proposed mechanisms are not necessarily mutually exclusive (Irving et al., 2021). While this model can explain viral replication in bats, it is inadequate to explain the eventual viral clearance. Furthermore, the similarity of innate immune responses to filoviruses in bat and human cell lines (Kuzmin et al., 2017) despite distinct clinical outcomes of the infections in bats versus humans contradicts these theories. Yet another study, based on genomic analysis, hypothesized that evolutionarily divergent genes could explain bat-human differences in responses to viruses (Pavlovich et al., 2018), which is similar to our approach; however, in our data, the genes highlighted in (Pavlovich et al., 2018) (NK receptors, MHC class I genes, and certain type I interferons) were not responsive to the infections.

Our study demonstrated that that the response of bats to filoviruses involves multiple interrelated processes, and the difference in the responses to infection between bats and humans is driven by genes that have evolutionarily diverged, altering the response of several pathways at the systemic level. This results in a measured inflammatory response that allows an adaptive response to be mounted to clear the infection and suggests potential therapeutic strategies for controlling the disease caused by filoviruses in humans.

## Methods

### Viruses

Recombinant wild-type MARV, strain 371Bat, was recovered similarly in BHK-21 cells (Albariño et al., 2013) and passaged twice in Vero E6 cells for amplification. Recombinant wild-type EBOV, strain Mayinga, was recovered from the full-length clone and support plasmids in HEK 293T (Whitfield et al., 2020) cells and passaged twice in Vero E6 cells for amplification.

### Bat experimental protocol

All animal procedures were performed in compliance with protocols approved by the Institutional Animal Care and Use Committee at the University of Texas Medical Branch at Galveston. Adult ERBs were obtained from Wildlife Safari (animals which were not infected) or CJG EXOTICS (animals which were subsequently infected) and quarantined for 30 days under ABSL-2 conditions. Animals were implanted with microchip transponders for animal ID and temperature data collection. For studies with MARV and EBOV, animals were transferred to the Galveston National Laboratory ABSL-4 facility. Animals were segregated into groups of three. In the infection study, except for one MARV-infected male (bat #3), all bats were female. Male bats are prone to fighting, and as such, should not be cohoused in non-flight caging. Each group was housed in a separate cage for inoculation with the same virus. After acclimation to the facility, animals were anesthetized with isoflurane and infected subcutaneously in the scapular region with 10^4^ PFU (titrated on Vero E6 cells) of MARV or EBOV. Every other day, animals were anesthetized by isoflurane, weighed, temperature was determined via transponder, and 100-150 µL of blood was collected from the propatagial vein. Blood was inactivated in 1 mL of TRIzol reagent (Thermo-Fisher Scientific). Samples were then removed from ABSL-4 containment, and RNA was extracted. ddRT-PCR with primers specific to the nucleoprotein (NP) gene was used to detect viremia. Animals were euthanized 48 hours after detection of MARV or EBOV RNA load under deep isoflurane sedation via cardiac exsanguination confirmed by bilateral open chest. Tissues were collected (listed in Table ST1) and immediately homogenized in an appropriate volume of TRIzol reagent and stored at −80°C. 1 cubic centimeter (cc) tissue sections were homogenized in minimal essential media (MEM) supplemented with 10% fetal bovine serum and stored at −80°C. Additional tissue sections were fixed in 10% neutral buffered formalin for histopathology. Tissues and PBMCs were also collected from three uninfected control animals.

### Leukocyte isolation

Leukocytes were isolated using ACK lysis buffer (Gibco). Ten volumes of lysis buffer were added to whole blood, incubated for 2-3 min, and then neutralized with complete DMEM medium containing 10% FBS. Following neutralization, samples were centrifuged at 250 *g* for 5 min at 4°C, after which the supernatant was decanted from the pellet. This process was repeated several times per sample until a white pellet of cells free of contaminating red blood cells remained. Because density gradient purification was not performed on these samples prior to or after red blood cell lysis, these leukocyte preparations were assumed to contain granulocytes in addition to PBMCs.

### mRNA sequencing

Total RNA was isolated from bat tissues using Ambion’s RNA isolation and purification kit. For most samples, polyA-tailed mRNA was selected using beads with oligo-deoxythymidine and then fragmented. A few samples with poor RIN (RNA Integrity Number) scores were treated with Ribominus (targeting human ribosomal RNA) to enrich for polyA-tailed mRNA before fragmentation. cDNA was synthesized using random hexamers and ligated with bar-coded adaptors compatible with Illumina’s NextSeq 500 sequencer. A total of 88 samples were sequenced on the NextSeq 500, as 75 base pair single reads.

### Bat mRNA sequence database

The extant bat genomes are not complete, and a comprehensive mRNA database does not exist. Thus, for this study, we constructed a custom non-redundant reference bat mRNA sequence database, which is available at the FiloBat website (Sachidanandam). We started with existing genome annotations (Jebb et al., 2019). The complications arising from splice variants were avoided by keeping only the longest transcript for each gene. We added missing annotations/sequences (e.g., CYP11B2 and PLG) to our database by assembling reads from our own sequence data. These required custom scripts as there often was not enough reads covering a transcript, which precluded the use of standard assembly tools. The gene sequences were collected from different bat species, so error-free reads might not map perfectly to the transcripts in the database. The database has sequences of 18,443 bat mRNAs and includes MARV and EBOV sequences. The gene sequences were collected from different bat species.

The genes were identified by homology to mouse and human genes, 16,004 bat genes had high similarity to human or mouse homologues, as defined by BLASTn with default settings identifying matches spanning the length of the mRNA. The set of remaining genes (2439) were labelled as divergent. Of these, 1,548 transcripts could be identified by increasing the sensitivity of BLASTn by reducing the word-size from 11 to 9, which is equivalent to matching at the protein level. Of the remaining 891 putative transcripts, homologues for 182 could be identified based on partial homology and domain structure, while the remainder (709 sequences whose names start with UNDEF) belonged to one of four classes, 1) aligned to un-annotated transcripts in the human genome, 2) non-coding RNAs, 3) transcripts unique to bats, or 4) assembly errors. We use capitalizations to represent bat gene symbols, as in the human gene nomenclature.

To identify genes within these divergent set that are relevant to our study, we then selected a subset of genes that had good expression (defined as tpm > 20) in at least one class of liver samples (MARV-infected, EBOV-infected, or uninfected) and responsive in either MARV-infected or EBOV-infected bat livers, which we defined as upregulated (log_2_ ratio > 0.6), or downregulated (log_2_ ratio < −0.6). We were left with 151 genes that are the foundation of our analyses of pathways involved in the response to filoviruses (Tables ST3-8).

### Expression Analyses

To determine transcript expression levels, we used Kallisto (Bray et al., 2016), because this tool uses pseudo-alignments and is relatively more tolerant of errors/variants in reads, which we expect here because the reads and mRNA sequences in the database do not always come from the same species. Kallisto uses a parameter “k” while indexing the database to specify how sensitive it is to matches with smaller k values leading to more spurious hits. We empirically determined k=23 to be an appropriate parameter value with which to index the reference mRNA dataset. We used tpm value as the transcript expression levels to determine changes in expression across samples.

We used viral transcripts to identify infected samples, which has previously helped us to identify and correct errors in annotation in some of the cell line data and also identified a problem with a published dataset (Hölzer et al., 2016), where all the naïve (uninfected) samples showed signs of viral infection. Furthermore, to ensure there was no mislabeling of tissue samples from different bats, we used single nucleotide variants in the sequenced reads to confirm that all tissue samples from an individual had the same set of variants.

Using clustering based on expression profiles and considering individual interferon responsive genes, it was clear that one non-infected control bat liver sample (labeled *cb1* in the FiloBat tool (Sachidanandam)) was reacting to some stimulus (injury or infection) compared to the other two control samples (*cb2* and *cb3* in the FiloBat tool (Sachidanandam)). Correlations between samples based on gene expressions profiles (Table ST9) shows that cb1 is an outlier and is closer in profile to the MARV-infected samples, suggesting an inflammatory response is present. This animal had an injury, and it is likely that inflammatory processes associated with this were responsible. Since we are interested in the innate response to infections, we had to exclude *cb1* from the controls and all analyses, but *cb1* data are available for exploration in the *filobat* tool. Most of our analyses concentrated on liver RNA transcripts since it had the strongest response, and the genes indicated that a variety of cell types were involved in the response, capturing the systemic nature of the response. Liver function impacts a wide range of systems involving inflammation, iron homeostasis, and blood pressure. Other organs, such as kidney and spleen provide additional support for what is observed in the liver. For some genes, we also used the transcriptional response in kidney (renin) and/or spleen (STING) to understand the regulation of pathways (e.g., renin is secreted by kidney and regulates the blood pressure system (Pickering, 1989)).

### Tools for data exploration and interrogation

To allow exploration of the data across various samples on a gene-by-gene basis, as well as analysis of viral expression in the samples, we developed a browser-based tool, *FiloBat*, using *Shiny* in R (Sachidanandam). Samples can also be compared using scatter plots and hierarchical clustering.

### Statistics

Large changes in expression profiles were readily detected by comparing averages across replicates, since such changes are less affected by noise; however, subtle changes (less than 2-fold) were difficult to reliably detect due to lack of power in the sample size and variability between samples and are mostly not considered. Since only two samples were left in the controls, we could not use the t-test to compute p-values for comparisons between the EBOV and control samples. Correlations between the various samples (Table ST9) suggest that we could perform comparisons between the MARV and the EBOV (which are broadly similar to the uninfected samples), and over 130 of the 194 genes highlighted in the study have FDR values below 0.25. Many of these (40 genes) would have exceeded this threshold if all the genes in the study were considered. The full table of genes with the p-values and FDR is available on the companion website (Sachidanandam).

### Pathway analyses

A fundamental assumption underlying our study is that bats are mammals that possess innate and adaptive responses to infections that roughly mirror those seen in humans. The data from comparative filovirus infections in human and bat cell lines supports this assumption (Kuzmin et al., 2017). To identify pathways of interest from particular genes, we used GO/pathways annotations of the human counterparts (The Gene Ontology Resource: 20 years and still GOing strong, 2019) and grouped them into functions that provided themes in the dataset. Using these themes, we identified other differentially expressed genes sharing these themes, identified by the GO annotations for human and mouse genes. This allowed us to build a picture of the pathways triggered by filovirus infections and delineate the ways in which the systemic bat responses differs from those seen in humans.

### Availability of data

All data underlying the balloon plots is available as csv files on the filobat website (Sachidanandam). Additionally, a fasta-format file, containing all the mRNA sequences used in our analysis, is also available on the website. The raw sequencing reads will be deposited with GEO, and the filobat site has several tools for analysis and exploration of data.

## Results

### Inoculation of bats with MARV and EBOV results in detectable viral replication in some organs

Nine ERBs were either inoculated subcutaneously with 10^4^ PFU of MARV or EBOV or were left uninfected (three in each group). Following inoculation, animals were observed at least daily and bled every other day. As the goal of the study was to investigate changes in gene expression in the early phase of the disease, MARV-infected animals were euthanized 48 hours after the expected peak of the viral replication based on a previously published study (Jones et al., 2015). Previous experimental infections of ERBs with EBOV resulted in detection of low levels of viral RNA (but not live virus) in the blood and some organs on days 3 – 16 after infection (Jones et al., 2015), suggesting a possible slow replication of the virus. Because of that, a later time point – day 11 – was selected for euthanasia of EBOV-infected ERBs. As expected, bats inoculated with MARV or EBOV showed no apparent clinical signs of disease or changes in behavior, with no significant effect on body weight and temperature (Fig. 1A, B). MARV or EBOV RNA was detected by droplet digital RT-PCR (ddRT-PCR) in the blood of infected bats (Fig. 1C). MARV was detected by plaque assay in livers and spleens of all inoculated animals and in the salivary glands of two animals and kidneys of one animal (Fig. 1D). By contrast, EBOV was detected in the livers of two inoculated animals and could not be reliably detected elsewhere (Fig. 1E).

**Fig. 1.**
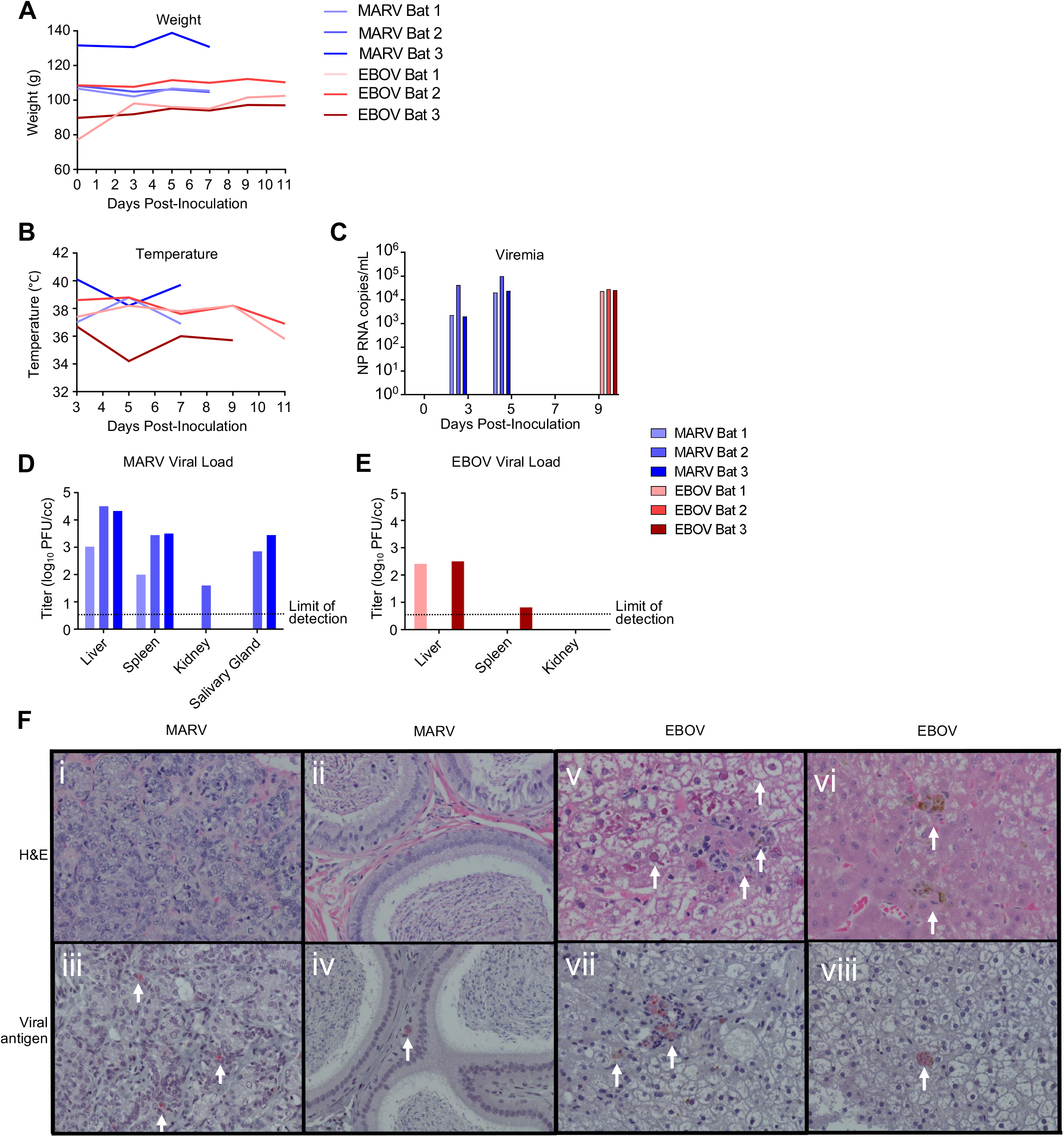
Infection of bats with filoviruses MARV and EBOV. **A – C.** Time course after infection for weight (A), temperature (B) and viral RNA specific for the NP gene in total RNA extracted from whole blood. Animals were euthanized 48 h after first viremic time point. **D**, **E.** Tissue viral loads on day 7 for EBOV and day 11 for MARV determined by plaque assay. Note that MARV bat 2 sensor failed. **F.** Histopathology and viral antigen in filovirus infected organs showing: (i) Lack of significant histopathological lesions in mammary tissue from MARV-infected bat 1, (ii) Lack of significant histopathological lesions within the interstitium of the epididymis from MARV-infected bat 3, (iii) IHC detection of MARV antigen within the interstitium of the glandular structures of the mammary gland from MARV-infected bat 1 (red pigment, arrows), (iv) IHC detection of MARV antigen within the interstitium of the epididymis from MARV-infected bat 3 (red pigment, arrows), (v) EBOV-infected bat 1 liver with marked histopathological changes, including cytoplasmic and nuclear inclusions (arrows), (vi) EBOV-infected bat 2 liver displaying a less dramatic presentation compared to bat 1 (arrows), (vii) IHC detection of EBOV antigen in the liver of EBOV-infected bat 1 (arrows), (viii) IHC detection of EBOV antigen in EBOV-infected bat 1 liver (an arrow).

No pathology was observed in sections of female mammary tissue or in the testes of MARV-inoculated animals (Fig. 1Fi,ii), that is consistent with previous reports (Jones et al., 2015). However, immunohistochemistry analysis demonstrated MARV antigen in both tissues, though this was focal in nature (Fig 1Fiii, iv), despite the absence of histopathological lesions in these organs. Two of the three EBOV-inoculated animals presented histopathological lesions in the liver, consisting of pigmented and unpigmented infiltrates of aggregated mononuclear cells compressing adjacent tissue structures, and eosinophilic nuclear and cytoplasmic inclusions (Fig. 1Fv,vi), changes consistent with previous reports (Krähling et al., 2010; Yuan et al., 2012). Immunohistochemical analysis demonstrated EBOV antigen in the liver of one animal, but very few foci were found (Fig 1Fvii, viii), suggesting limited viral replication. Overall, the histological studies demonstrated moderate amounts of MARV antigen and low amounts of EBOV antigen, that is consistent with the virus load data generated by plaque titration and ddRT-PCR, and very limited pathological changes.

### MARV and EBOV infection affects the transcriptome of multiple organs

To examine the transcriptional response to filovirus infections, we performed deep sequencing of mRNA from liver, spleen, kidneys, lungs and large intestine collected from filovirus inoculated and uninfected bats (Methods, Table ST1). We focused on analysis of the transcriptional response in liver, spleen and kidneys as these organs are classic targets of filovirus infections. Consistent with prior reports that liver is the primary target of MARV (Becker et al., 1995), and with our findings (Fig. 1), MARV transcripts were the most abundant in liver (79 transcripts-per-million or tpm) and also present in spleen (56 tpm), intestine (10 tpm) and lungs (2 tpm) but not present in kidneys (Table ST2). EBOV transcripts were detected at very low levels (< 1 tpm) in the livers of inoculated bats and were not detectable in other tissues.

Remarkably, although the highest levels of viral transcripts were detected in the liver, gene expression patterns were altered in all three tissues subjected to transcriptome analysis, liver, spleen, and kidneys, involving thousands of genes, suggesting a systemic response (Fig. 2, S1). The changes were higher in the livers of bats infected with MARV than EBOV (Fig. 2), which can be explained by the much more abundant replication of MARV (Fig. 1). The sets of genes affected by the two infections were not completely identical, suggesting some responses are virus-specific (Fig. 2, S1). The differences in the patterns of gene expression could also arise from different levels of MARV and EBOV replication.

**Fig. 2.**
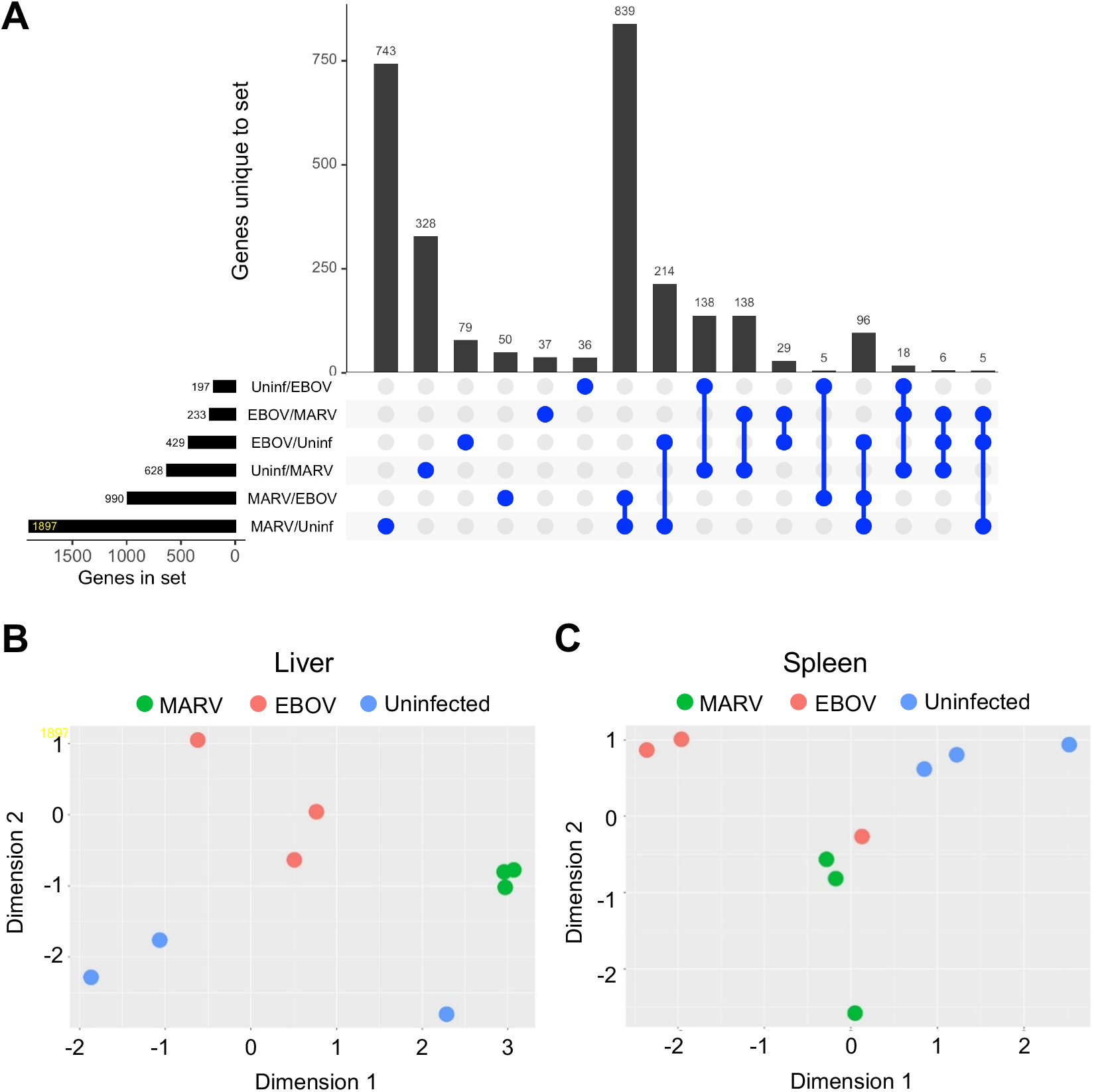
Changes in gene expression patterns in bats after infections with MARV and EBOV: **A.** Upset plot, uninfected bat livers. EBOV: EBOV-infected bat livers, MARV: MARV-infected bat livers. Each row in the lower panel represents a set with the corresponding bars at the lower left representing membership in the sets. There are six sets of genes, e.g., EBOV/Uninf comprises of genes at least 2-fold upregulated in EBOV infection, compared to the uninfected samples, while Uninf/EBOV is genes at least 2-fold down regulated in EBOV samples compared to the uninfected samples. The vertical blue lines with bulbs represent sets of intersections with the main bar plot (top) is number of genes unique to that intersection, so the total belonging to the set Uninf/EBOV, as an example, is a sum of the numbers in all sets that have Uninf/EBOV as a member (41+203+6+31=281). The last bar with 6 members is the set of genes common to EBOV/MARV, EBOV/Uninf and MARV/Uninf. Many more genes responded to MARV infection than to EBOV infection (1,897 genes in MARV/Uninf versus 429 genes in EBOV/Uninf). The much greater number of MARV-specific (MARV/EBOV) genes than EBOV-specific (EBOV/MARV) genes is most likely due to the different levels of infections; when the samples were collected, EBOV was cleared, while MARV was in the process of being cleared. Some differences between MARV and EBOV are likely due to interferon antagonism differentially mediated by VP40 and VP35 in MARV and VP40 and VP24 in EBOV. 2 **B, C.** Multidimensional scaling (MDS) plots of the merged gene expression data from livers of MARV-infected, EBOV-infected and uninfected bats. There is a clear separation between MARV infections, EBOV infections and the uninfected sample both in livers (**B**) and spleens (**C**). Virus-specific signatures were also detected in kidneys (**Fig. S1**), implying the response to filovirus infections extends to the whole animal. The location of one blue dot in panel A is different from the other two samples from uninfected bats, as also seen in the correlations shown in **Table ST9**. The different pattern for this bat may be explained by its reaction to some stimulus, either an infection or an injury; the animal has been excluded from the downstream analysis.

### Understanding the response to filovirus infection using the framework of evolutionarily divergent bat genes

The stark difference in the outcomes of filovirus infections in bats versus humans, despite substantial parts of the response being similar, suggests that evolutionarily divergent genes cause some pathways to have different outcomes. We identified divergent genes in the bat genome and used this restricted set to identify pathways of interest. We then used all genes in these pathways for further analysis. Amongst the bat genes, 2,439 genes had diverged from their human or mouse homologues according to our criteria (Fig. 3, Methods).

**Fig. 3.**
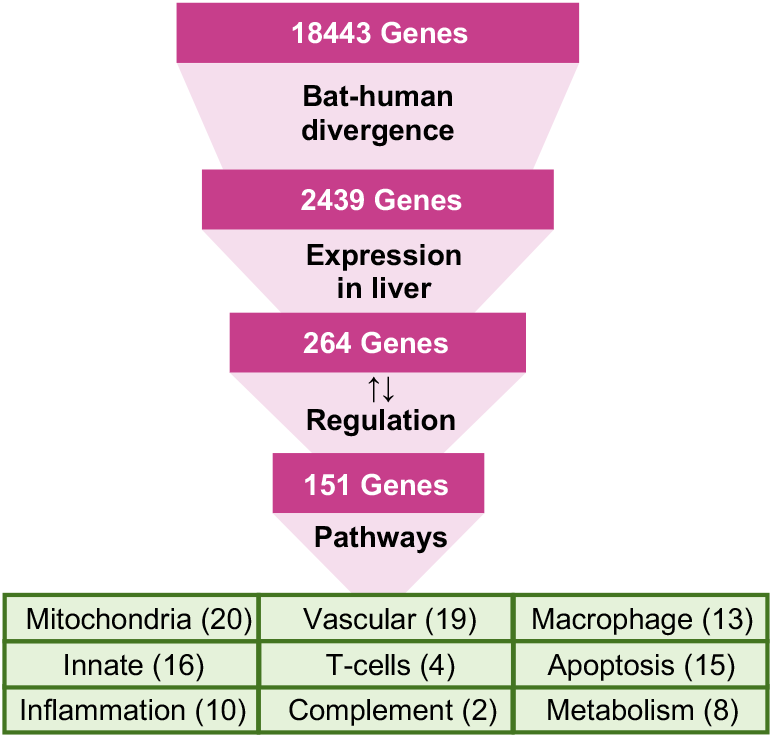
Analysis of evolutionary divergent genes as an approach for understanding the response to filovirus infections. The approach used in this study was based on identification of pathways relevant to the bat’s resilience to filoviral infections. It was assumed that the most homologous genes perform similar functions in bats and humans, and bat genes evolutionarily divergent from their human homologs have a greater probability of having altered functions. The divergent genes were identified using homology detected through BLAST alignments of the corresponding sequences from the two species. Of these, genes differentially expressed in livers of infected bats were identified. The pathways they influence were explored to evaluate the systemic response to filovirus infections in bats and to identify key differences from the responses in humans. Blood pressure, coagulation and iron homeostasis pathways were altered most prominently. The analysis demonstrated changes in glycolysis, which is controlled by hypoxia, which shift the balance of macrophage activation from the M1 (proinflammatory) to the M2 (anti-inflammatory) state. These changes create a pro-inflammatory state that modulate the response and allows the adaptive immune system to clear the infection early, and anti-inflammatory response late. The identified pathways demonstrated incomplete activation of the complement system, likely compromising the antibody response, but strong activation of T cell response which is likely to play a major role in clearing the infection. The identified pathways are interconnected, as shown in Fig. 6.

The major benefit of limiting the genes to the divergent set is that we ameliorate the multiple-testing problem. For example, 130 liver-specific genes were differentially responsive upon comparing the response to MARV and EBOV infections with a false discovery rate (FDR) < 0.25. In contrast, starting from the full list of bat genes in the genome would yield only 90 genes with FDR < 0.25, due to the false positives from the larger set of background genes reducing the signal (table shown in accompanying website (Sachidanandam)).

Of the 2,439 divergent genes (Fig. 3), only 264 were expressed at more than 20 tpm in at least one set of liver samples (MARV, EBOV, or uninfected) and only 151 of these genes were responsive to either MARV or EBOV (Fig. S2). These 151 genes were the basis of the first step of pathway analysis (Tables ST3-8). The most abundant group in this set comprised genes related to mitochondria (20 genes), followed by genes involved in the vascular system (19), innate immunity (16), tissue regeneration and apoptosis (15), macrophages (13), inflammation (10), metabolism and fatty-acid oxidation (8), T cells (4), complement system (2), digestion (5), and toxin processing (3).

Based on this preliminary list of pathways, we focused on the *entire* transcriptomes of the following systems: 1) innate immune system (Fig. S3) which includes the interferon stimulated genes (Fig. S4), most of which are conserved between bats and humans, phagocytosis by macrophages, natural killer cells and the complement system; 2) inflammatory response, including acute phase proteins, macrophages functions involving metabolism, fatty-acid oxidation, mitochondrial abundance and function, and tissue regeneration and apoptosis; and 3) blood-related physiological systems, involving the regulation of blood pressure, coagulation, and iron homeostasis.

### Effects of filovirus infections on the transcriptional response

#### MARV and EBOV infection induce initial inflammation, evidenced primarily by an acute phase response

Acute phase proteins (APP) are produced by hepatocytes in the liver in response to inflammatory cytokines, such as interleukin (IL)-1, (IL-6) and tumor necrosis factor *α* (TNF*α*) and are an important part of the innate immune response (Kushner, 1982; Gabay and Kushner, 1999). Upon inflammation, the concentration of positive APPs, including serum amyloid A1 (SAA1) and serum amyloid A2 (SAA2), increase dramatically (> 10-fold) in the serum (Gauldie et al., 1987), while the concentration of negative APPs, including transferrin (TF) and albumin (ALB), decreases (Moshage et al., 1987).

We found that MARV, and to a lesser extent EBOV, infection induced an APP response in liver, spleen, and kidney, with the largest changes in APP expression (>10-fold) observed in the liver (Table 1, Fig. 4, S2A, S2B). SAA1 and SAA2 expression increased to a similar degree in all tissues, including tissues in which the viruses were not detected. At the same time, we detected no expression of C-reactive protein (CRP), an APP used as a marker for inflammation/acute-phase-response in humans (Table 1, Fig. 4), likely because it is not expressed in bats. We draw this conclusion in part because we were also unable to identify evidence of a CRP response in an analysis of public mRNA-seq data from infected samples from various species of bats (data not shown). Consistent with the induction of SAA1 and SAA2, we also detected induction of other markers of inflammation including orosomucoid 2 (ORM2), ceruloplasmin (CP), hepcidin (HAMP) and the microsomal glutathione S-transferases (MGST1, MGST2) (Dvash et al., 2015) (Table 1, Fig. 4).

**Fig. 4.**
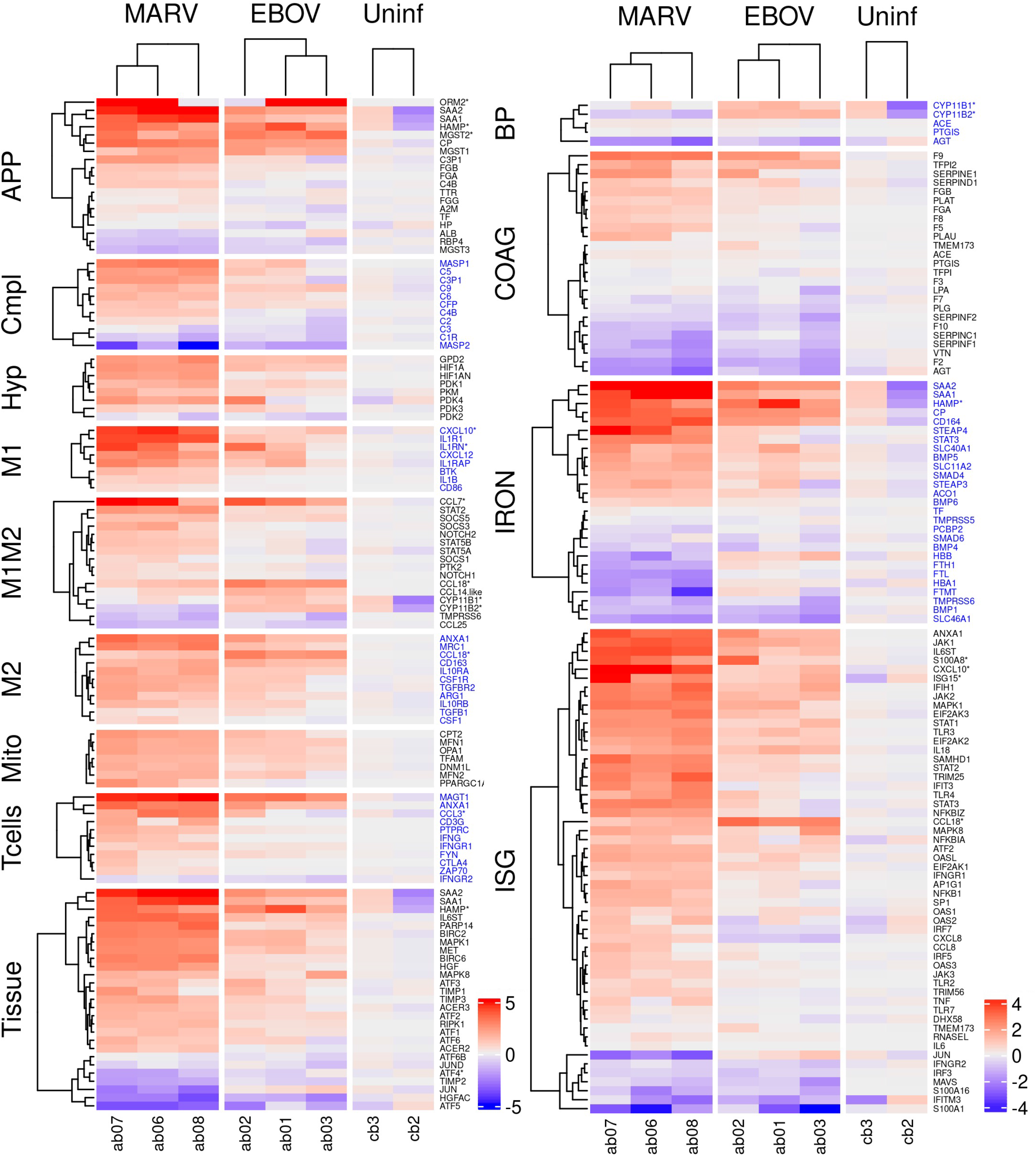
Differential expression of genes belonging to the indicated pathways by MARV and EBOV infections in livers of bats. The **c**olumns show differentially expressed genes from livers of three MARV-infected bats, three EBOV-infected bats and two uninfected bats. The values are log_2_ of the fpm values, with the mean values from uninfected animals subtracted. Broadly, the majority of the genes are in a quiescent, low expression state in the uninfected samples, and get stimulated upon infections, with greater effects in the case of MARV and intermediate effects in case of EBOV. APP: acute phase response proteins; Compl: complement; Hyp: hypoxia related genes; M1 and M2: genes specific to M1 or M2 macrophages, respectively; M1M2: genes common to M1 and M2 macrophages; Mito: genes expressed in mitochondria; Tcells: genes expressed in T cells; Tissue: genes involved in tissue regeneration and apoptosis; BP: genes involved in regulation of blood pressure; ISG: interferon stimulated genes; IRON: genes involved in iron homeostasis; COAG: genes involved in coagulation. A **\*** after a gene name indicates that the bat version is divergent from its human counterpart. Alternate blocks of gene names are colored black/blue to allow easy visual distinction of the blocks. **Fig. S2A** and **S2B** show corresponding figures for kidneys and spleens.

**Table 1.** Acute phase proteins in livers respond strongly to inflammatory cytokines. Basal expression levels in tpm units are shown in the Uninfected column. The fold change upon infection is shown in the MARV and EBOV columns. SAA1/2, CP were highly expressed in livers normally and also get upregulated by filovirus infections, with a greater upregulation by MARV. ORM2, MGST2 are upregulated, but from a low basal expression level. CRP, used as a marker for acute phase response in humans, is highlighted in gray to emphasize that it does not appear to be expressed in this bat species and might be absent in bats altogether. TF was highly expressed in all samples, but the expression did not change in response to filoviral infections, while TTR was not expressed in any of the samples. A similar inflammation was detected in the liver upon both MARV and EBOV infection, despite the lack of viral transcripts in the liver of EBOV infected animals. *Bat genes divergent from their respective human homologs.

| **Positive APPs** | **MARV**<br>**(fold change)** | **EBOV**<br>**(fold change)** | **Uninfected**<br>**(tpm)** |
| --- | --- | --- | --- |
| Serum Amyloid A 1 (SAA1) | 21X | 3X | 6858 |
| Serum Amyloid A 2 (SAA2) | 39X | 5X | 440 |
| Ceruloplasmin (CP) | 10X | 5X | 129 |
| HAMP* | 8.5X | 10X | 211 |
| Orosomucoid 2 Alpha1-Acid glycoprotein (ORM2*) | 34X | 47X | 14 |
| Microsomal Glutathione S-Transferase MGST1 | 4X | 4X | 277 |
| MGST2* | 11X | 16X | 5.5 |
| MGST3 | 0.4X | 0.7X | 461 |
| Fibrinogen (FGA) | 2X | 1X | 1277 |
| Fibrinogen (FGB) | 2X | 1X | 9007 |
| Fibrinogen (FGG) | 1X | 1X | 6070 |
| C4B | 2X | 1X | 1015 |
| C3P1 | 6X | 1X | 31 |
| Haptoglobin (HP) | 1.1X | 0.7X | 15906 |
| Alpha2-Macroglobulin (A2M) | 1.3X | 1X | 409 |
| C-reactive protein (CRP) | N/A | N/A | N/A |
| **Negative APPs** |  |  |  |
| Albumin (ALB) | 0.6X | 1.2X | 51400 |
| Transferrin (TF) | 1X | 1X | 22856 |
| Transthyretin (TTR) | 2X | 2X | 1 |
| Retinol Binding protein (RBP4) | 0.5X | 0.6X | 3107 |

#### Expression of key components of the classical complement pathway is inhibited by filovirus infection

The complement system, a part of the immune system, has three pathways: the classical pathway, the mannose-binding lectin pathway and the alternative pathway (Noris and Remuzzi, 2013). The classical pathway recognizes antibodies bound to antigens, the lectin pathway is triggered by the binding of lectin to sugar molecules on pathogens, and the alternative pathway is triggered by C3 recognition of a hydroxyl or amino group of a molecule on a pathogen surface.

Several key genes associated with the complement pathway were upregulated by filovirus infection, including complement component 3 precursor pseudogene (C3P1), complement C4B (C4B), complement C5 (C5), complement C9 (C9), complement C6 (C6), and mannan binding lectin serine peptidase 1 (MASP1), while others were downregulated or not expressed, including complement C1r (C1R), complement C3 (C3), complement C8 gamma chain (C8G), and mannan binding lectin serine peptidase 2 (MASP2) (Fig. 4, S5A). These observations indicated that the complement pathway is affected by filovirus infection and suggests that some protective mechanisms involving the complement system such as antibody-dependent complement deposition may be compromised. Of note, most of the genes implicated in this pathway are non-divergent between bats and humans.

#### Infected bats exhibit transcriptional signatures of T cell activity

CD4*^+^* T cells recognize peptides presented on MHC class II molecules expressed by antigen presenting cells (APCs), while CD8^+^ T cells recognize peptides presented by MHC class I molecules, expressed by all nucleated cells (Germain, 2002). CD8^+^ T cells are cytotoxic and can kill virus-infected cells. Multiple genes expressed only by CD8^+^ T cells, including C-C motif chemokine ligand 3 (CCL3), annexin A1 (ANAX1), T cell immunoglobulin and mucin domain containing 4 (TIMD4) and magnesium transporter 1 (MAGT1), were upregulated in the liver by filovirus infection (Fig. 4, S5B), indicating an induction of T cell response.

#### MARV and EBOV infections affect blood related physiological systems

Blood carries nutrients, oxygen, as well as the cells and molecules that constitute the immune response and inflammation. The key physiological systems connected to the immune response and inflammation are iron homeostasis, blood pressure, and blood coagulation. We found that expression of genes involved in regulation of all three of these systems was affected by MARV and EBOV infections, as detailed below.

#### Genes regulating iron homeostasis

The absorption and availability of iron, an essential component of heme needed for oxygen transport, is tightly regulated (Knutson, 2017). In humans, most iron in the body is located in hemoglobin (66%) and myoglobin (10%) (Fraction of iron in body that is present in h - Human Homo sapiens - BNID 104009), while the remainder is stored mostly in macrophages in the liver, which take up iron through the CD163 receptor. Iron is exported from macrophages and absorbed from food (Prentice et al., 2012) through ferroportin (SLC40A1/FPN1).

MARV and EBOV infections changed the expression of multiple genes involved in iron homeostasis. Hepcidin (HAMP), which controls iron homeostasis by binding ferroportin (Kohgo et al., 2008; Przybyszewska and Żekanowska, 2014) thereby causing its degradation as well as blocking the export of iron, was induced in infected livers (Fig. 4, S6, Table 1). Similarly, ceruloplasmin (CP) (an APP, Table 1), which enables the formation of the transferrin-iron complex and is also involved in processing copper (Wessling-Resnick, 2018), was also induced. In the cytosol, iron is bound to ferritin (comprised of a heavy chain, FTH1 and a light chain FTL), synthesized by cells in response to increased iron (Kohgo et al., 2008). In mitochondria, iron is bound to the mitochondrial ferritin (FTMT) (Gao and Chang, 2014). Both FTH1 and FTMT were downregulated in MARV-infected but not EBOV-infected bats (Fig. 4, S6); the difference can be explained by higher MARV replication relative to EBOV. Furthermore, MARV infection was associated with lowered hemoglobin (HBB) expression, suggesting impairment of red blood cell production. Consistent with this conclusion, CD164, which suppresses hematopoietic cell proliferation, was also upregulated by MARV, and to a lesser degree, EBOV infection (Fig. 4, S6). Thus, hematopoiesis may be impaired in MARV-infected ERBs, but not in EBOV-infected ERBs, the difference consistent with the levels of viral replication for the respective viruses in ERBs. Furthermore, as the higher levels of HAMP in EBOV-infected bats do not result in lower levels of FTH1/FTMT, the regulation of iron by HAMP in bats is likely to be diverged from the homologous process in humans (Kohgo et al., 2008; Przybyszewska and Żekanowska, 2014).

#### Genes regulating blood pressure

The primary means of blood pressure regulation is renal expression of renin, which converts angiotensinogen (AGT) to angiotensin I. Angiotensin converting enzyme (ACE) converts angiotensin I to angiotensin II, which constricts blood vessels to increase blood pressure (Lu et al., 2016). Both MARV and EBOV infections downregulated AGT, resulting in depletion of the substrate for ACE, limiting the potential for blood pressure to increase even with upregulation of ACE (Fig. 4, S7). Another mechanism regulating blood pressure involves cytochrome P450 family 11 subfamily B member 2 (CYP11B2), which increases aldosterone levels that increases blood volume and consequently, the blood pressure (Ji et al., 2015). CYP11B2 was found to be down regulated in MARV infected bats in this study (Fig. 4), further suggesting the possibility that low blood pressure is a response to MARV infection. In the case of EBOV, CYP11B2 was not downregulated (Fig. 4), possibly due to the low level of viral replication. However, blood pressure was not measured in the bats in this study.

#### Genes regulating in blood coagulation

Mechanisms that control blood pressure also impact coagulation and vice versa, increasing coagulation leads to higher blood pressure through constriction of blood vessels. For example, fibrinogen B (FGB) is cleaved by thrombin to generate fibrin (which forms the clots), and cleavage products of FGB promote vasoconstriction. The complement pathway also impacts coagulation (Conway, 2018).

Coagulation or clotting is a complex process involving a cascade of activation reactions that finally results in thrombin forming clots (Fig. S8). Since many of the gene products involved need to be activated, information on the coagulation state is not readily observed from mRNA abundance, which can at most indicate the protein abundance, not their states of activation. Of the genes in the coagulation pathway, MARV and EBOV infections upregulated one set of genes: coagulation factor IX (F9), tissue factor pathway inhibitor 2 (TFPI2), serpin family E member 1 (SERPINE1), SERPIND1, fibrinogen beta chain (FGB), plasminogen activator tissue type (PLAT), and downregulated another set: thrombin (F2), vitronectin (VTN), SERPINF1, SERPINC1, coagulation factor X (F10) and plasminogen (PLG) (Fig. 4, S8, S9). There were other genes which were not strongly up or down regulated upon the infections.

In the coagulation cascade (Fig. S8), tissue factor III (encoded by the F3 gene) activates coagulation factor VII (encoded by the F7 gene), which then activates several factors in the cascade, eventually activating coagulation factor X (encoded by the F10 gene) which in turn activates coagulation factor II (encoded by the F2 gene) to form thrombin, which enables fibrin synthesis, leading to clot formation. Thrombin is also involved in a positive feedback enabling more thrombin to be created. An opposing process involves the conversion of plasminogen to plasmin (facilitated by the plasminogen activators uPA and tPA) which promotes fibrinolysis (Hoover-Plow, 2010), leading to dissolution of clots. SERPINE1 inhibits this process by inhibiting the activity of uPA and tPA.

F3 was not abundant in any sample, and F2 was downregulated in response to MARV and EBOV infections, suggesting thrombin production was curtailed; SERPINE1 was upregulated, which can only prevent dissolution of clots, but does not enhance coagulation directly. Angiotensin II, which increases blood pressure, also increases thrombin formation and impairs fibrinolysis (Miesbach, 2020). AGT, the precursor of angiotensin II, was downregulated by MARV and EBOV infections, which is expected to lower angiotensin II levels and concomitantly decrease blood pressure and coagulation. Interestingly, a link between some acute human viral infections and hypotension have been established including for COVID-19 (Kariyanna et al., 2020). Thus, we conclude that filovirus infections of bats lead to reduced coagulation. Together, these events are likely to reduce the effects of inflammation on the vascular system.

#### MARV and EBOV infection leads to an early transition from proinflammatory M1 to anti-inflammatory M2-dominated populations of macrophages

Macrophages recognize and phagocytize microbes and damaged host cells, as a part of the innate immune response, and are an important early target for filoviruses (Becker et al., 1995). Macrophages can be in a continuum between the M1 state, an inflammatory state enabling apoptosis, and the M2 state, an anti-inflammatory state assisting tissue regeneration (Fig. S5C). A key difference between the M1 and M2 states lies in their metabolism; the M1 state is characterized by hypoxia and glycolysis metabolism (Wang et al., 2017), while the M2 state is characterized by fatty acid metabolism and elevated mitochondrial activity (Mills and O’Neill, 2016).

We have found that key markers of the M1 state were upregulated in livers of infected bats and more so in MARV infected animals. These markers included interferon regulatory factor 5 (IRF5), nuclear factor kappa B subunit 1 (NFKB1), adaptor related protein complex 1 subunit gamma 1 (AP1G1), signal transducer and activator of transcription 1 (STAT1), and suppressor of cytokine signaling 3 (SOCS3) (Fig. 4, S10, S11). Likewise, hypoxia inducible factor 1 subunit alpha (HIF1A) (Nizet and Johnson, 2009; Krawczyk et al., 2010; Tannahill et al., 2013), which promotes mitophagy and glycolysis metabolism to induce M1 polarization, was also upregulated in infected livers, again more so in MARV infection. Pyruvate kinase M1/2 (PKM1/2), which activate HIF1A, and pyruvate dehydrogenase kinase 1 (PDK), which enhances M2 polarization in response to hypoxia (Tan et al., 2015), were also upregulated. Again, upregulation occurred to a greater degree in MARV than EBOV (Fig. 4, S10, S11).

Markers of the M2 state, such as mannose receptor C-Type 1 (MRC1), arginase 1(ARG1), IL-10 and transforming growth factor beta 1 (TGFB1) (Ferrante et al., 2013), were highly expressed in livers of bats infected with both viruses (Fig. 4, S10, S11F), suggesting that M2 macrophages were also present. Several genes related to fatty acid oxidation in M2 macrophages were upregulated by MARV and EBOV infections (Tables ST3A, ST5, ST7). Particularly, we observed upregulation of carnitine palmitoyl transferase 2 (CPT2), a gene associated with fatty acid transport; the effect was greater in MARV infection. Infected bats also exhibited upregulation of multiple markers of mitochondria abundance, another characteristic of M2 macrophages (Fig. S11C). These included transcription factor A mitochondrial (TFAM), OPA1 mitochondrial dynamin like GTPase (OPA1), mitofusin (MFN1/2), and dynamin 1 like (DNM1L). Also upregulated upon MARV infection are genes involved in mitochondrial biogenesis (Imamura and Matsumoto, 2017), hepatocyte growth factor/tyrosine kinase MET (HGF/MET) and peroxisome proliferator-activated receptor gamma coactivator 1 alpha (PPARGC1A).

Prolonged M1 activity can be harmful to tissues due to their induction of inflammation and apoptosis. This activity is modulated by a negative feedback system that shifts macrophages from the M1 state to the M2 state (Atri et al., 2018; Parisi et al., 2018), controlling inflammation during infection and facilitating the transition to tissue repair and regeneration (Helming, 2011; Jenkins et al., 2011). In our data, the transcriptomes of the MARV-infected liver samples exhibited markers for M1 and M2 macrophages, while in the EBOV-infected liver samples, markers for M2 macrophages dominated. These data suggest that the presence of both M1 and M2 macrophages upon MARV infection might be indicative of the greater magnitude and duration of its replication, and the presence of M2 macrophages upon EBOV infection indicates at a conversion from M1 to M2 state during or after the virus clearance.

Consistent with this conversion, we observed changes in cellular energy metabolism that are associated with the M1 to M2 transition. The mitochondrial glycerol-3-phosphate dehydrogenase (GPD2), identified as a contributor to the shift in core macrophage metabolism associated with the M1 to M2 transition during infection (Murphy, 2019), was found to be upregulated by filovirus infections (Fig. 5, S11B). Filovirus infections of bats resulted in upregulation of hypoxia inducible factor 1 subunit alpha inhibitor (HIF1AN), the inhibitor of HIF1A; inactivating HIF1A also promotes M2 polarization (Werno et al., 2010), suggesting a mechanism of M2 polarization in filovirus infected bats. Interestingly, in MARV-infected bats, which demonstrated a mixed M1-M2 response, we observed downregulation of ferritin (FTH1/FTL), which is synthesized in response to iron, and HBB (a reflection of hemoglobin levels), and upregulation of HAMP, which reduces the levels of iron. In EBOV-infected animals, which demonstrated a more pronounced M2 response (Fig. S11D, S11E), no downregulation of FTH1/FTL, a small upregulation of HBB and a stronger upregulation of HAMP was detected. These data are consistent with the data that an increased availability of iron promotes the M1 to M2 polarization shift (Agoro et al., 2018). These lines of evidence further support our findings of a polarization bias toward M2 during filovirus infections.

**Fig. 5.**
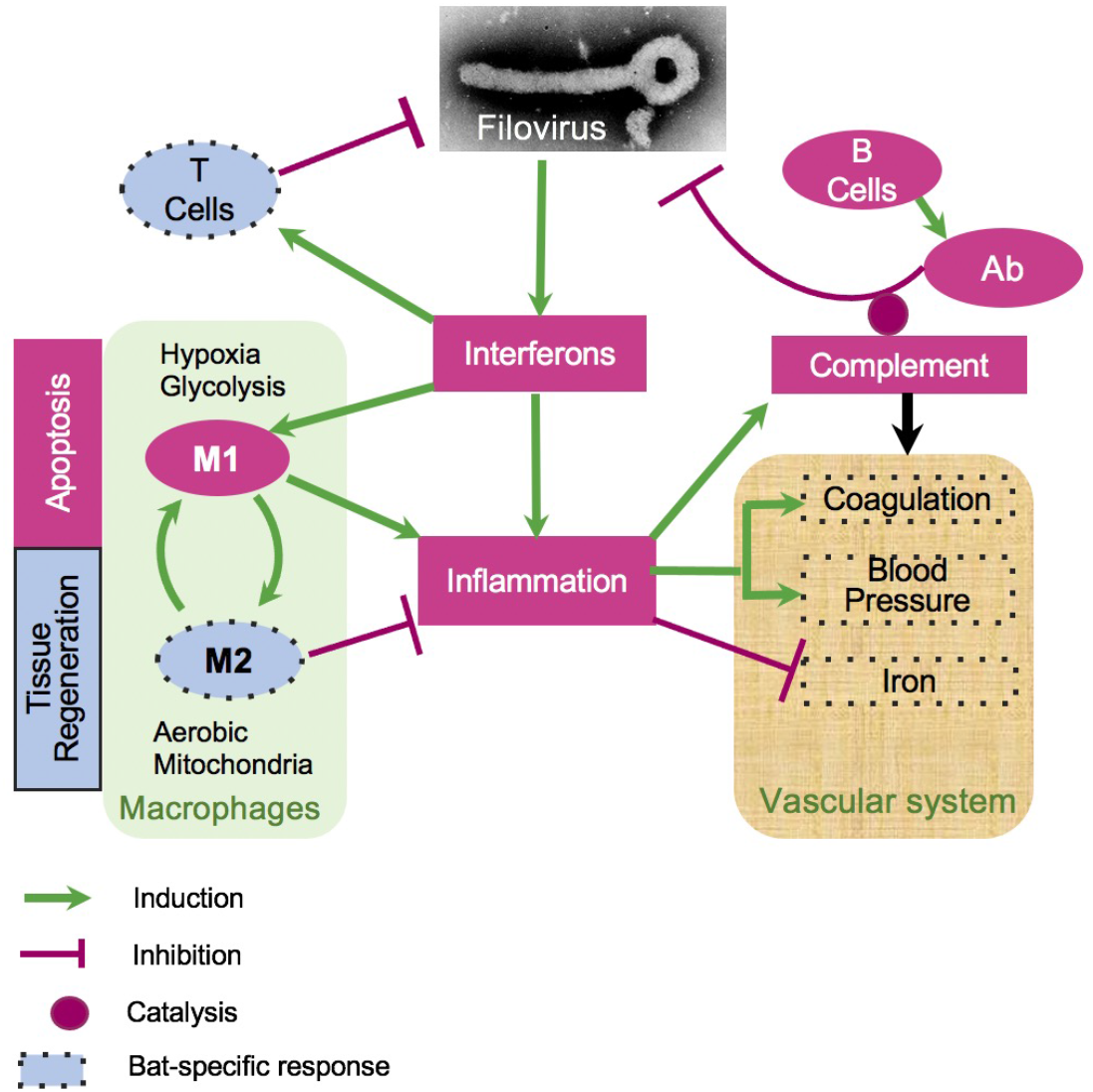
A model of bat response to filovirus infections. Interferon stimulated genes (ISG, Fig. 5,S2,S3) cause inflammation, which triggers the acute phase response (APR, Table 1, Fig. 5), leading to a cascade of reactions affecting regulation of HAMP (iron, Fig. 5,S6), coagulation (Fig. 5, S7, S8), blood pressure (Fig. 5, S7) and stimulating M1 macrophages (Fig. 5, S5, S10, S11). The pro-inflammatory M1 macrophages phagocytose infected cells and promote apoptosis. Over the course of infection, M1 macrophages get converted to anti-inflammatory M2 macrophages. This process is accompanied by activation of fatty acid oxidation and mitochondrial activity, which are the hallmarks of the M2 macrophage responses (Fig. 5, S10, S11). On the other hand, activation of the complement system is incomplete, potentially leading to a reduced antibody activity (Fig. 5, S5B). Furthermore, filovirus infections were accompanied by downregulation of blood pressure (Fig. 5, S7) and the coagulation system (Fig. 5, S9). The infections are also accompanied by shift in genes regulating iron homeostasis (Fig. 5,S7, S8, S9, S11), which are generally consistent with a greater activation of the M2 macrophage response in case of EBOV infections. The infections resulted in activation of CD8^+^ T cell response (Fig. S5C), presumably contributing to clearance of the infections. The dotted boundaries indicate at pathways which responded to filovirus infections in bats differently from human.

## Discussion

Over the past few years, multiple high-impact pathogens associated with bats have emerged or re-emerged, the most notable of which are MARV, EBOV, SARS-CoV and SARS-CoV-2. As a result, the role of bats as reservoirs for a diverse array of viruses and their ability to tolerate viral infections that cause severe disease in humans has become a topic of considerable interest. Several hypotheses, mostly centered on the innate immune system, have been proposed to explain this unique aspect of bat biology. In these hypotheses, various aspects of bat innate immunity are either more or less potent than their human counterparts. One of these hypotheses posits that some bat species constitutively express interferons, leading to a basal level of innate immune activation (Glennon et al., 2015; Zhou et al., 2016; Pavlovich et al., 2018). However, in humans persistent interferon expression leads to lowered resistance to infections due to dysregulation of iron homeostasis (Rossi and Deepe, 2020) and are associated with various pathologies (Ng et al., 2019). Patients with trisomy have elevated constitutive expression of interferons and exhibit weaker immune response in COVID-19 (Sullivan et al., 2016; Wadm et al., 2020). These pathologies and prior work with filoviruses demonstrating that the innate response in bat cells is robust, and similar to that observed in human cell lines (Kuzmin et al., 2017) suggests that interferons in bats are induced similarly to that observed in humans. Thus, the hypothesis of constitutive high expression of interferons in bats is inconsistent with our data. Another hypothesis suggests that components of the innate immune response, e.g., stimulator of interferon response cGAMP interactor 1 (STING/TMEM173), are less effective in bats, allowing viruses to survive in the host (Xie et al., 2018). This mechanism cannot explain the virus clearance, although it might explain the lack of symptoms during the infections in bats. The eventual clearance of filoviral infections by bats suggests a more complex process involving both the innate and adaptive immune systems.

In this study, all MARV-inoculated bats were productively infected, and our virology and histopathology data in MARV-infected bats are consistent with previous reports, including viral replication in the mammary glands and testes (Jones et al., 2015). Unexpectedly, evidence of productive, while limited, infection was identified in two of the three EBOV inoculated animals, although the detection of virus in the livers by plaque assay and immunohistochemistry identified only a small number of foci in the liver of one animal (Fig. 1E). These data contrast the prior reports (Jones et al., 2015; Paweska et al., 2016), and suggests that ERBs may not be truly refractory to EBOV infection. However, given the low titers detected, and the limited nature of the observed immunostaining, it remains to be determined whether the virus can be maintained in this bat species in nature. To reveal molecular mechanisms that underlie the resistance of bats to the disease caused by MARV and EBOV, we developed a model of bat response to filovirus infections (Fig. 5). Our analysis shows that the majority of interferon response genes are not divergent from their human homologs, consistent with our prior observations that the innate responses to filoviruses are quite similar in human and bat cells (Kuzmin et al., 2017). These structural and functional similarities suggest that the ability of bats to tolerate infections with multiple viral pathogens without a disease is unlikely to be explained by features of the interferon response. These data are in agreement with the recently published study on MARV infection in ERBs (Guito et al., 2021).

A key feature of filovirus infection in humans is an inflammatory response leading to the expression of APPs and stimulation of M1 macrophages. In humans, a major APP protein is CRP, which binds to microorganisms, including viruses, assists in complement binding to foreign and damaged cells, and enhances phagocytosis by macrophages (opsonin-mediated phagocytosis) (Wu et al., 2015). We found no evidence of CRP expression in bats by mRNA-seq data, while other APPs were conserved and are listed in Table 1. It is possible the lack of CRP response contributes to a lowered inflammation, explaining the resilience of bats to infections by many viruses. As a counterargument, we must keep in mind that in mice, CRP is not an acute phase protein (Du Clos, 2003).

We found evidence in the response of complement genes that an effector component of the antibody response may be weakened by incomplete complement activation. This is consistent with the previous reports that antibody-mediated virus neutralization is not the dominant mechanism of filovirus clearance in ERBs (Schuh et al., 2019). The robust CD8^+^ T cell activity implied by our mRNA-seq data suggests that control and clearance of filovirus infection in bats may instead depend upon a robust T cell response. This is consistent with what is known in humans, where individuals who recover from filovirus infections tend to mount robust T cell responses (McElroy et al., 2015; Dahlke et al., 2017; Reynard et al., 2019), and have higher levels of CD40L expression, a marker for T cell activity (Reynard et al., 2019), although recovery of humans from filovirus infections also correlates with induction of the antibody response (Ksiazek et al., 1999).

The macrophage response was one of the more notable points of divergence between the human response to filovirus infection and what we observed in infected bats. We identified markers of both M1 and M2 macrophages in ERBs infected with MARV, suggesting that macrophage populations in the animals were in the process of transition towards the anti-inflammatory M2 state, associated with tissue repair and regeneration as opposed to the classic pro-inflammatory M1 state. In particular, the modulation of the innate response facilitated by M2 macrophages is important for T cell mediated clearance of viruses (Atri et al., 2018). In EBOV-infected animals, where viral replication was far more limited, our data indicate that the macrophage population was further along in the transition to M2 polarization by the time of euthanasia, likely due to the viral replication being successfully controlled. The generalized anti-inflammatory state observed in bats during filovirus infection, especially the early transition towards M2 macrophage polarization, may suggest a way to prevent the immunopathology associated with filovirus infections in humans, including cytokine storm and DIC. Supporting this, an mRNA-seq study conducted with PBMCs isolated from EBOV-infected humans found that individuals who succumbed to disease showed stronger upregulation of interferon signaling and acute phase response-related genes than survivors during the acute phase of infection (Liu et al., 2017), suggesting that a moderated innate response improves outcomes in filovirus infection. Furthermore, pharmacological inhibition of toll-like receptor 4 signaling promotes survival in an animal model of filovirus infections (Younan et al., 2017).

Our data suggest that the bat vascular response to infection differs from that in humans. Humans infected with MARV or EBOV frequently present with hemorrhagic manifestations and dysregulated coagulation in the form of DIC (Dovih et al., 2019). We identified transcriptional patterns consistent with vasodilation and reduced potential for coagulation which could result in a state of low blood pressure, and reduced coagulation. This state may be protective, as it might be expected to prevent DIC. These findings are consistent with results from a study in humans infected with EBOV which analyzed 55 biomarkers in blood and found that viremia was associated with elevated levels of tissue factor and tissue plasminogen activator, consistent with coagulopathy (McElroy et al., 2014).

Our results suggest that reducing the hyperinflammatory response (Younan et al., 2017) or controlling coagulopathy (Rasmussen et al., 2014) in humans during filovirus infection may have a therapeutic benefit by preventing damage to the host and allowing other processes to clear the infection. This could be achieved by inhibiting IL-6 signaling either by targeting the cytokine (by Clazakizumab, Olokizumab, Sirukumab or Siltuximab), or its cognate receptor (by Tocilizumab or Sarilumab) or its trans-signaling by blocking the soluble receptor (sIL-6R) (by Olamkicept) (Jones and Hunter, 2021). Strikingly, a recent COVID-19 study demonstrated tocilizumab reduced the likelihood of progression to the composite outcome of mechanical ventilation or death (Salama et al., 2021).

We demonstrated that in bats, filovirus infections upregulates MGST1 and MGST2 (Table 1), which induce leukotrienes (LTC4) and prostaglandin E, both of which induce inflammation (Dvash et al., 2015). This is a potential druggable target, as these molecules are targeted by several therapeutic agents. Thus, inflammation caused by filovirus infections could also be potentially targeted by another class of anti-inflammatory agents such as LTC4 inhibitors, used to treat asthma.

Our results also suggest that upon filovirus infections bats may naturally vasodilate and reduce their blood pressure (mimicking the action of ACE inhibitors), while the endothelial system becomes anti-thrombotic. This suggestion is consistent with the results of the field trials of ACE inhibitors and statins in human Ebola virus disease that have already demonstrated some success (Fedson, 2015). The potential involvement of vasodilation also suggests that prostaglandin I_2_ (PGI_2_, known as the drug Epoprostenol), a powerful vasodilator and anti-coagulant that acts by binding to the prostacyclin receptor (Cacione et al., 2016), could be investigated in human filovirus infections as a means of emulating the physiological conditions (low blood pressure and coagulation) that our data suggest may have protective effects.

In humans, high levels of HAMP cause iron to be sequestered in the liver, reducing levels of iron in blood (lower ferritin). Our observations indicate that in EBOV infected bats high HAMP expression is decoupled from the levels of iron, as both ferritin and HAMP are induced. Thus, HAMP inhibitors, which are used to treat anemia, might recreate in humans the state seen in bats under filoviral infection.

The changes in gene expression patterns that we have observed in infected bats suggest that the interconnected pathways regulating coagulation, vasodilation, iron homeostasis, inflammation, the interferon response, and the adaptive response contribute to the unique response of bats to filovirus infection. This response appears to be avoiding immunopathology by tempering of the inflammatory response to infection. In particular, the anti-inflammatory state (macrophages in the M2 state) and the altered state of the blood-related physiological systems seem to be important in preventing pathology and facilitating the ultimate clearance of the viruses. The filobat website (Sachidanandam), developed to accompany this paper, allows easy exploration of our data and comparisons with future studies that might be performed. The unexpected features of bat responses to filovirus infections may aid in the development of new strategies to effectively mitigate and treat the disease caused by filovirus infections in humans.

## Acknowledgements

Oliver Fregoso provided early comments and encouragement for our approach. Yelena Ginzburg helped in clarifying the role of HAMP in iron homeostasis. Radhika Patnala and Arne Fabritius of Sci-Illustrate helped in the generation of figures. Yuri Lazebnik provided immense help by editing the text and offering critical comments that clarified our message. Viviana Simon read early versions and gave suggestions and encouragement. This work was primarily funded by the grant HDTRA1-16-1-0033 from the Defense Threat Reduction Agency (CFB, AB, RS). Work of RS was also partially supported by the grant R01-AI136916 from NIH/NIAID.

---

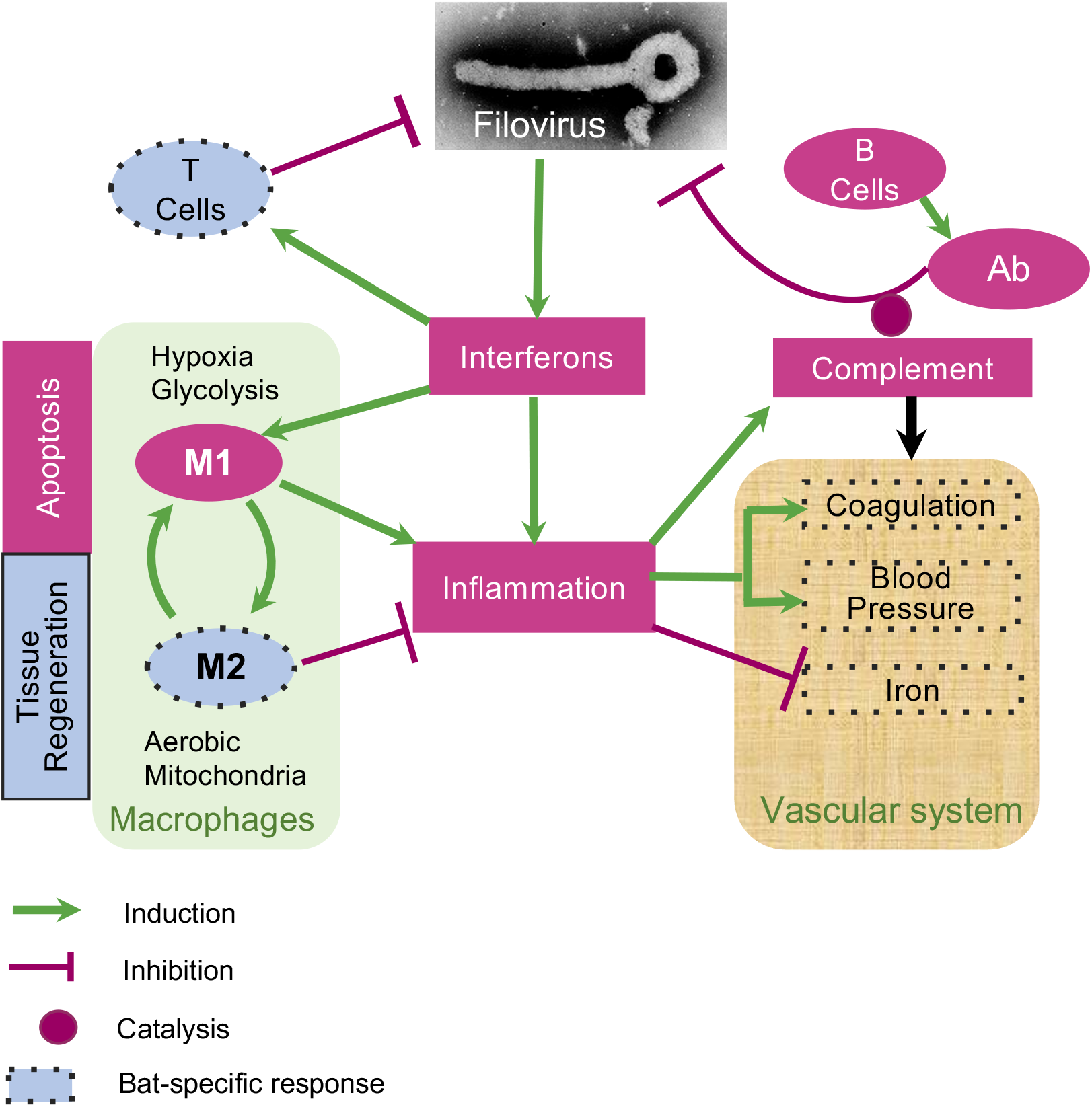

## REFERENCES

Agoro, R., Taleb, M., Quesniaux, V. F. J., and Mura, C. (2018). Cell iron status influences macrophage polarization. PLOS ONE 13, e0196921. doi:10.1371/journal.pone.0196921.

Albariño, C. G., Uebelhoer, L. S., Vincent, J. P., Khristova, M. L., Chakrabarti, A. K., McElroy, A., et al. (2013). Development of a reverse genetics system to generate recombinant Marburg virus derived from a bat isolate. Virology 446, 230–237. doi:10.1016/j.virol.2013.07.038.

Amman, B. R., Bird, B. H., Bakarr, I. A., Bangura, J., Schuh, A. J., Johnny, J., et al. (2020). Isolation of Angola-like Marburg virus from Egyptian rousette bats from West Africa. Nat Commun 11, 510. doi:10.1038/s41467-020-14327-8.

Amman, B. R., Carroll, S. A., Reed, Z. D., Sealy, T. K., Balinandi, S., Swanepoel, R., et al. (2012). Seasonal Pulses of Marburg Virus Circulation in Juvenile Rousettus aegyptiacus Bats Coincide with Periods of Increased Risk of Human Infection. PLOS Pathogens 8, e1002877. doi:10.1371/journal.ppat.1002877.

Amman, B. R., Jones, M. E. B., Sealy, T. K., Uebelhoer, L. S., Schuh, A. J., Bird, B. H., et al. (2015). Oral shedding of Marburg virus in experimentally infected Egyptian fruit bats (Rousettus aegyptiacus). J. Wildl. Dis. 51, 113–124. doi:10.7589/2014-08-198.

Atri, C., Guerfali, F. Z., and Laouini, D. (2018). Role of Human Macrophage Polarization in Inflammation during Infectious Diseases. Int J Mol Sci 19. doi:10.3390/ijms19061801.

Becker, S., Spiess, M., and Klenk, H. D. (1995). The asialoglycoprotein receptor is a potential liver-specific receptor for Marburg virus. J. Gen. Virol. 76 (Pt 2), 393–399. doi:10.1099/0022-1317-76-2-393.

Bray, N. L., Pimentel, H., Melsted, P., and Pachter, L. (2016). Near-optimal probabilistic RNA-seq quantification. Nature Biotechnology 34, 525–527. doi:10.1038/nbt.3519.

Cacione, D. G., Macedo, C. R., and Baptista-Silva, J. C. (2016). Pharmacological treatment for Buerger’s disease. Cochrane Database of Systematic Reviews. doi:10.1002/14651858.CD011033.pub3.

Conway, E. M. (2018). Complement-coagulation connections. Blood Coagulation & Fibrinolysis 29, 243. doi:10.1097/MBC.0000000000000720.

Dahlke, C., Lunemann, S., Kasonta, R., Kreuels, B., Schmiedel, S., Ly, M. L., et al. (2017). Comprehensive Characterization of Cellular Immune Responses Following Ebola Virus Infection. J. Infect. Dis. 215, 287–292. doi:10.1093/infdis/jiw508.

Dhama, K., Karthik, K., Khandia, R., Chakraborty, S., Munjal, A., Latheef, S. K., et al. (2018). Advances in Designing and Developing Vaccines, Drugs, and Therapies to Counter Ebola Virus. Front Immunol 9, 1803. doi:10.3389/fimmu.2018.01803.

Dovih, P., Laing, E. D., Chen, Y., Low, D. H. W., Ansil, B. R., Yang, X., et al. (2019). Filovirus-reactive antibodies in humans and bats in Northeast India imply zoonotic spillover. PLOS Neglected Tropical Diseases 13, e0007733. doi:10.1371/journal.pntd.0007733.

Du Clos, T. W. (2003). C-reactive protein as a regulator of autoimmunity and inflammation. Arthritis Rheum. 48, 1475–1477. doi:10.1002/art.11025.

Dvash, E., Har-Tal, M., Barak, S., Meir, O., and Rubinstein, M. (2015). Leukotriene C_4_ is the major trigger of stress-induced oxidative DNA damage. Nature Communications 6, 10112. doi:10.1038/ncomms10112.

Fedson, D. S. (2015). A Practical Treatment for Patients With Ebola Virus Disease. J Infect Dis 211, 661–662. doi:10.1093/infdis/jiu474.

Ferrante, C. J., Pinhal-Enfield, G., Elson, G., Cronstein, B. N., Hasko, G., Outram, S., et al. (2013). The adenosine-dependent angiogenic switch of macrophages to an M2-like phenotype is independent of interleukin-4 receptor alpha (IL-4Rα) signaling. Inflammation 36, 921–931. doi:10.1007/s10753-013-9621-3.

Fraction of iron in body that is present in h - Human Homo sapiens - BNID 104009 Available at: https://bionumbers.hms.harvard.edu/bionumber.aspx?s=n&v=3&id=104009 [Accessed March 15, 2021].

Gabay, C., and Kushner, I. (1999). Acute-phase proteins and other systemic responses to inflammation. N. Engl. J. Med. 340, 448–454. doi:10.1056/NEJM199902113400607.

Gao, G., and Chang, Y.-Z. (2014). Mitochondrial ferritin in the regulation of brain iron homeostasis and neurodegenerative diseases. Front Pharmacol 5. doi:10.3389/fphar.2014.00019.

Gauldie, J., Richards, C., Harnish, D., Lansdorp, P., and Baumann, H. (1987). Interferon beta 2/B-cell stimulatory factor type 2 shares identity with monocyte-derived hepatocyte-stimulating factor and regulates the major acute phase protein response in liver cells. Proc. Natl. Acad. Sci. U.S.A. 84, 7251– 7255. doi:10.1073/pnas.84.20.7251.

Geisbert, T. W., Young, H. A., Jahrling, P. B., Davis, K. J., Larsen, T., Kagan, E., et al. (2003). Pathogenesis of Ebola Hemorrhagic Fever in Primate Models. Am J Pathol 163, 2371–2382.

Germain, R. N. (2002). T-cell development and the CD4–CD8 lineage decision. Nat Rev Immunol 2, 309–322. doi:10.1038/nri798.

Glennon, N. B., Jabado, O., Lo, M. K., and Shaw, M. L. (2015). Transcriptome Profiling of the Virus-Induced Innate Immune Response in Pteropus vampyrus and Its Attenuation by Nipah Virus Interferon Antagonist Functions. J. Virol. 89, 7550–7566. doi:10.1128/JVI.00302-15.

Guito, J. C., Prescott, J. B., Arnold, C. E., Amman, B. R., Schuh, A. J., Spengler, J. R., et al. (2021). Asymptomatic Infection of Marburg Virus Reservoir Bats Is Explained by a Strategy of Immunoprotective Disease Tolerance. Curr Biol 31, 257–270.e5. doi:10.1016/j.cub.2020.10.015.

Helming, L. (2011). Inflammation: Cell Recruitment versus Local Proliferation. Current Biology 21, R548– R550. doi:10.1016/j.cub.2011.06.005.

Hölzer, M., Krähling, V., Amman, F., Barth, E., Bernhart, S. H., Carmelo, V. A. O., et al. (2016). Differential transcriptional responses to Ebola and Marburg virus infection in bat and human cells. Scientific Reports 6, 34589. doi:10.1038/srep34589.

Hoover-Plow, J. (2010). Does plasmin have anticoagulant activity? Vasc Health Risk Manag 6, 199–205. doi:10.2147/vhrm.s9358.

Imamura, R., and Matsumoto, K. (2017). Hepatocyte growth factor in physiology and infectious diseases. Cytokine 98, 97–106. doi:10.1016/j.cyto.2016.12.025.

Irving, A. T., Ahn, M., Goh, G., Anderson, D. E., and Wang, L.-F. (2021). Lessons from the host defences of bats, a unique viral reservoir. Nature 589, 363–370. doi:10.1038/s41586-020-03128-0.

Jebb, D., Huang, Z., Pippel, M., Hughes, G. M., Lavrichenko, K., Devanna, P., et al. (2019). Six new reference-quality bat genomes illuminate the molecular basis and evolution of bat adaptations. bioRxiv, 836874. doi:10.1101/836874.

Jenkins, S. J., Ruckerl, D., Cook, P. C., Jones, L. H., Finkelman, F. D., van Rooijen, N., et al. (2011). Local macrophage proliferation, rather than recruitment from the blood, is a signature of TH2 inflammation. Science 332, 1284–1288. doi:10.1126/science.1204351.

Ji, X., Qi, H., Li, D.-B., Liu, R.-K., Zheng, Y., Chen, H.-L., et al. (2015). Associations between human aldosterone synthase CYP11B2 (−344T/C) gene polymorphism and antihypertensive response to valsartan in Chinese patients with essential hypertension. Int J Clin Exp Med 8, 1173–1177.

Jones, M. E. B., Amman, B. R., Sealy, T. K., Uebelhoer, L. S., Schuh, A. J., Flietstra, T., et al. (2019). Clinical, Histopathologic, and Immunohistochemical Characterization of Experimental Marburg Virus Infection in A Natural Reservoir Host, the Egyptian Rousette Bat (Rousettus aegyptiacus). Viruses 11. doi:10.3390/v11030214.

Jones, M. E. B., Schuh, A. J., Amman, B. R., Sealy, T. K., Zaki, S. R., Nichol, S. T., et al. (2015). Experimental Inoculation of Egyptian Rousette Bats (Rousettus aegyptiacus) with Viruses of the Ebolavirus and Marburgvirus Genera. Viruses 7, 3420–3442. doi:10.3390/v7072779.

Jones, S. A., and Hunter, C. A. (2021). Is IL-6 a key cytokine target for therapy in COVID-19? Nature Reviews Immunology, 1–3. doi:10.1038/s41577-021-00553-8.

Kariyanna, P. T., Sutarjono, B., Grewal, E., Singh, K. P., Aurora, L., Smith, L., et al. (2020). A Systematic Review of COVID-19 and Myocarditis. Am J Med Case Rep 8, 299–305.

Knutson, M. D. (2017). Iron transport proteins: Gateways of cellular and systemic iron homeostasis. J. Biol. Chem. 292, 12735–12743. doi:10.1074/jbc.R117.786632.

Kohgo, Y., Ikuta, K., Ohtake, T., Torimoto, Y., and Kato, J. (2008). Body iron metabolism and pathophysiology of iron overload. Int J Hematol 88, 7–15. doi:10.1007/s12185-008-0120-5.

Kortepeter, M. G., Dierberg, K., Shenoy, E. S., Cieslak, T. J., and Medical Countermeasures Working Group of the National Ebola Training and Education Center’s (NETEC) Special Pathogens Research Network (SPRN) (2020). Marburg virus disease: A summary for clinicians. Int J Infect Dis 99, 233–242. doi:10.1016/j.ijid.2020.07.042.

Krähling, V., Dolnik, O., Kolesnikova, L., Schmidt-Chanasit, J., Jordan, I., Sandig, V., et al. (2010). Establishment of fruit bat cells (Rousettus aegyptiacus) as a model system for the investigation of filoviral infection. PLoS Negl Trop Dis 4, e802. doi:10.1371/journal.pntd.0000802.

Krawczyk, C. M., Holowka, T., Sun, J., Blagih, J., Amiel, E., DeBerardinis, R. J., et al. (2010). Toll-like receptor-induced changes in glycolytic metabolism regulate dendritic cell activation. Blood 115, 4742– 4749. doi:10.1182/blood-2009-10-249540.

Ksiazek, T. G., Rollin, P. E., Williams, A. J., Bressler, D. S., Martin, M. L., Swanepoel, R., et al. (1999). Clinical virology of Ebola hemorrhagic fever (EHF): virus, virus antigen, and IgG and IgM antibody findings among EHF patients in Kikwit, Democratic Republic of the Congo, 1995. J Infect Dis 179 Suppl 1, S177-187. doi:10.1086/514321.

Kushner, I. (1982). The phenomenon of the acute phase response. Ann. N. Y. Acad. Sci. 389, 39–48. doi:10.1111/j.1749-6632.1982.tb22124.x.

Kushner, I. Acute phase reactants. UpToDate. Available at: https://www.uptodate.com/contents/acute-phase-reactants.

Kuzmin, I. V., Schwarz, T. M., Ilinykh, P. A., Jordan, I., Ksiazek, T. G., Sachidanandam, R., et al. (2017). Innate Immune Response of Bat and Human Cells to Filoviruses: Commonalities and Distinctions. J. Virol.

Leroy, E. M., Epelboin, A., Mondonge, V., Pourrut, X., Gonzalez, J.-P., Muyembe-Tamfum, J.-J., et al. (2009). Human Ebola outbreak resulting from direct exposure to fruit bats in Luebo, Democratic Republic of Congo, 2007. Vector Borne Zoonotic Dis. 9, 723–728. doi:10.1089/vbz.2008.0167.

Liu, X., Speranza, E., Muñoz-Fontela, C., Haldenby, S., Rickett, N. Y., Garcia-Dorival, I., et al. (2017). Transcriptomic signatures differentiate survival from fatal outcomes in humans infected with Ebola virus. Genome Biology 18, 4. doi:10.1186/s13059-016-1137-3.

Lu, H., Cassis, L. A., Kooi, C. W. V., and Daugherty, A. (2016). Structure and functions of angiotensinogen. Hypertens Res 39, 492–500. doi:10.1038/hr.2016.17.

Marí Saéz, A., Weiss, S., Nowak, K., Lapeyre, V., Zimmermann, F., Düx, A., et al. (2015). Investigating the zoonotic origin of the West African Ebola epidemic. EMBO Mol Med 7, 17–23. doi:10.15252/emmm.201404792.

McElroy, A. K., Akondy, R. S., Davis, C. W., Ellebedy, A. H., Mehta, A. K., Kraft, C. S., et al. (2015). Human Ebola virus infection results in substantial immune activation. Proc. Natl. Acad. Sci. U.S.A. 112, 4719– 4724. doi:10.1073/pnas.1502619112.

McElroy, A. K., Erickson, B. R., Flietstra, T. D., Rollin, P. E., Nichol, S. T., Towner, J. S., et al. (2014). Ebola hemorrhagic Fever: novel biomarker correlates of clinical outcome. J. Infect. Dis. 210, 558–566. doi:10.1093/infdis/jiu088.

Miesbach, W. (2020). Pathological Role of Angiotensin II in Severe COVID-19. TH Open 4, e138–e144. doi:10.1055/s-0040-1713678.

Mills, E. L., and O’Neill, L. A. (2016). Reprogramming mitochondrial metabolism in macrophages as an anti-inflammatory signal. Eur. J. Immunol. 46, 13–21. doi:10.1002/eji.201445427.

Moshage, H. J., Janssen, J. A., Franssen, J. H., Hafkenscheid, J. C., and Yap, S. H. (1987). Study of the molecular mechanism of decreased liver synthesis of albumin in inflammation. J. Clin. Invest. 79, 1635– 1641. doi:10.1172/JCI113000.

Murphy, M. P. (2019). Rerouting metabolism to activate macrophages. Nat Immunol, 1–3. doi:10.1038/s41590-019-0455-5.

Ng, C. S., Kato, H., and Fujita, T. (2019). Fueling Type I Interferonopathies: Regulation and Function of Type I Interferon Antiviral Responses. Journal of Interferon & Cytokine Research 39, 383–392. doi:10.1089/jir.2019.0037.

Nizet, V., and Johnson, R. S. (2009). Interdependence of hypoxic and innate immune responses. Nat. Rev. Immunol. 9, 609–617. doi:10.1038/nri2607.

Noris, M., and Remuzzi, G. (2013). Overview of Complement Activation and Regulation. Semin Nephrol 33, 479–492. doi:10.1016/j.semnephrol.2013.08.001.

Olival, K. J., Islam, A., Yu, M., Anthony, S. J., Epstein, J. H., Khan, S. A., et al. (2013). Ebola virus antibodies in fruit bats, bangladesh. Emerging Infect. Dis. 19, 270–273. doi:10.3201/eid1902.120524.

Parisi, L., Gini, E., Baci, D., Tremolati, M., Fanuli, M., Bassani, B., et al. (2018). Macrophage Polarization in Chronic Inflammatory Diseases: Killers or Builders? J Immunol Res 2018. doi:10.1155/2018/8917804.

Pavlovich, S. S., Lovett, S. P., Koroleva, G., Guito, J. C., Arnold, C. E., Nagle, E. R., et al. (2018). The Egyptian Rousette Genome Reveals Unexpected Features of Bat Antiviral Immunity. Cell 173, 1098–1110.e18. doi:10.1016/j.cell.2018.03.070.

Paweska, J. T., Jansen van Vuren, P., Fenton, K. A., Graves, K., Grobbelaar, A. A., Moolla, N., et al. (2015). Lack of Marburg Virus Transmission From Experimentally Infected to Susceptible In-Contact Egyptian Fruit Bats. J. Infect. Dis. 212 Suppl 2, S109–118. doi:10.1093/infdis/jiv132.

Paweska, J. T., Jansen van Vuren, P., Masumu, J., Leman, P. A., Grobbelaar, A. A., Birkhead, M., et al. (2012). Virological and Serological Findings in Rousettus aegyptiacus Experimentally Inoculated with Vero Cells-Adapted Hogan Strain of Marburg Virus. PLoS One 7. doi:10.1371/journal.pone.0045479.

Paweska, J. T., Storm, N., Grobbelaar, A. A., Markotter, W., Kemp, A., and Jansen van Vuren, P. (2016). Experimental Inoculation of Egyptian Fruit Bats (Rousettus aegyptiacus) with Ebola Virus. Viruses 8. doi:10.3390/v8020029.

Pickering, T. G. (1989). Renovascular hypertension: etiology and pathophysiology. Semin Nucl Med 19, 79–88. doi:10.1016/s0001-2998(89)80003-0.

Pourrut, X., Souris, M., Towner, J. S., Rollin, P. E., Nichol, S. T., Gonzalez, J.-P., et al. (2009). Large serological survey showing cocirculation of Ebola and Marburg viruses in Gabonese bat populations, and a high seroprevalence of both viruses in Rousettus aegyptiacus. BMC Infect. Dis. 9, 159. doi:10.1186/1471-2334-9-159.

Prentice, A. M., Doherty, C. P., Abrams, S. A., Cox, S. E., Atkinson, S. H., Verhoef, H., et al. (2012). Hepcidin is the major predictor of erythrocyte iron incorporation in anemic African children. Blood 119, 1922– 1928. doi:10.1182/blood-2011-11-391219.

Przybyszewska, J., and Żekanowska, E. (2014). The role of hepcidin, ferroportin, HCP1, and DMT1 protein in iron absorption in the human digestive tract. Prz Gastroenterol 9, 208–213. doi:10.5114/pg.2014.45102.

Rasmussen, A. L., Okumura, A., Ferris, M. T., Green, R., Feldmann, F., Kelly, S. M., et al. (2014). Host genetic diversity enables Ebola hemorrhagic fever pathogenesis and resistance. Science 346, 987–991. doi:10.1126/science.1259595.

Reynard, S., Journeaux, A., Gloaguen, E., Schaeffer, J., Varet, H., Pietrosemoli, N., et al. (2019). Immune parameters and outcomes during Ebola virus disease. JCI Insight 4. doi:10.1172/jci.insight.125106.

Rossi, D. C. P., and Deepe, G. S. (2020). Type I Interferons Rule with an Iron Fist. Cell Host & Microbe 27, 317–319. doi:10.1016/j.chom.2020.02.007.

Rougeron, V., Feldmann, H., Grard, G., Becker, S., and Leroy, E. M. (2015). Ebola and Marburg haemorrhagic fever. J. Clin. Virol. 64, 111–119. doi:10.1016/j.jcv.2015.01.014.

Sachidanandam, R. FiloBat website. FiloBat website. Available at: https://katahdin.girihlet.com/shiny/bat/ [Accessed May 9, 2019].

Salama, C., Han, J., Yau, L., Reiss, W. G., Kramer, B., Neidhart, J. D., et al. (2021). Tocilizumab in Patients Hospitalized with Covid-19 Pneumonia. New England Journal of Medicine 384, 20–30. doi:10.1056/NEJMoa2030340.

Schuh, A. J., Amman, B. R., Jones, M. E. B., Sealy, T. K., Uebelhoer, L. S., Spengler, J. R., et al. (2017a). Modelling filovirus maintenance in nature by experimental transmission of Marburg virus between Egyptian rousette bats. Nat Commun 8, 14446. doi:10.1038/ncomms14446.

Schuh, A. J., Amman, B. R., Sealy, T. K., Kainulainen, M. H., Chakrabarti, A. K., Guerrero, L. W., et al. (2019). Antibody-Mediated Virus Neutralization Is Not a Universal Mechanism of Marburg, Ebola, or Sosuga Virus Clearance in Egyptian Rousette Bats. J. Infect. Dis. 219, 1716–1721. doi:10.1093/infdis/jiy733.

Schuh, A. J., Amman, B. R., Sealy, T. K., Spengler, J. R., Nichol, S. T., and Towner, J. S. (2017b). Egyptian rousette bats maintain long-term protective immunity against Marburg virus infection despite diminished antibody levels. Sci Rep 7, 8763. doi:10.1038/s41598-017-07824-2.

Sullivan, K. D., Lewis, H. C., Hill, A. A., Pandey, A., Jackson, L. P., Cabral, J. M., et al. (2016). Trisomy 21 consistently activates the interferon response. Elife 5. doi:10.7554/eLife.16220.

Tan, Z., Xie, N., Cui, H., Moellering, D. R., Abraham, E., Thannickal, V. J., et al. (2015). Pyruvate dehydrogenase kinase 1 participates in macrophage polarization via regulating glucose metabolism. J. Immunol. 194, 6082–6089. doi:10.4049/jimmunol.1402469.

Tannahill, G. M., Curtis, A. M., Adamik, J., Palsson-McDermott, E. M., McGettrick, A. F., Goel, G., et al. (2013). Succinate is an inflammatory signal that induces IL-1β through HIF-1α. Nature 496, 238–242. doi:10.1038/nature11986.

The Gene Ontology Resource: 20 years and still GOing strong (2019). Nucleic Acids Res 47, D330–D338. doi:10.1093/nar/gky1055.

Towner, J. S., Amman, B. R., Sealy, T. K., Carroll, S. A. R., Comer, J. A., Kemp, A., et al. (2009). Isolation of genetically diverse Marburg viruses from Egyptian fruit bats. PLoS Pathog 5, e1000536. doi:10.1371/journal.ppat.1000536.

Wadm, M., Ec. 15, 2020, and Pm, 5:00 (2020). COVID-19 is 10 times deadlier for people with Down syndrome, raising calls for early vaccination. Science | AAAS. Available at: https://www.sciencemag.org/news/2020/12/covid-19-10-times-deadlier-people-down-syndrome-raising-calls-early-vaccination [Accessed December 17, 2020].

Wang, T., Liu, H., Lian, G., Zhang, S.-Y., Wang, X., and Jiang, C. (2017). HIF1α-Induced Glycolysis Metabolism Is Essential to the Activation of Inflammatory Macrophages. Mediators Inflamm. 2017, 9029327. doi:10.1155/2017/9029327.

Werno, C., Menrad, H., Weigert, A., Dehne, N., Goerdt, S., Schledzewski, K., et al. (2010). Knockout of HIF-1α in tumor-associated macrophages enhances M2 polarization and attenuates their pro-angiogenic responses. Carcinogenesis 31, 1863–1872. doi:10.1093/carcin/bgq088.

Wessling-Resnick, M. (2018). Crossing the Iron Gate: Why and How Transferrin Receptors Mediate Viral Entry. Annu. Rev. Nutr. 38, 431–458. doi:10.1146/annurev-nutr-082117-051749.

Whitfield, Z. J., Prasad, A. N., Ronk, A. J., Kuzmin, I. V., Ilinykh, P. A., Andino, R., et al. (2020). Species-Specific Evolution of Ebola Virus during Replication in Human and Bat Cells. Cell Rep 32, 108028. doi:10.1016/j.celrep.2020.108028.

Wu, Y., Potempa, L. A., El Kebir, D., and Filep, J. G. (2015). C-reactive protein and inflammation: conformational changes affect function. Biol. Chem. 396, 1181–1197. doi:10.1515/hsz-2015-0149.

Xie, J., Li, Y., Shen, X., Goh, G., Zhu, Y., Cui, J., et al. (2018). Dampened STING-Dependent Interferon Activation in Bats. Cell Host Microbe 23, 297–301.e4. doi:10.1016/j.chom.2018.01.006.

Younan, P., Ramanathan, P., Graber, J., Gusovsky, F., and Bukreyev, A. (2017). The Toll-Like Receptor 4 Antagonist Eritoran Protects Mice from Lethal Filovirus Challenge. MBio 8. doi:10.1128/mBio.00226-17.

Yuan, J., Zhang, Y., Li, J., Zhang, Y., Wang, L.-F., and Shi, Z. (2012). Serological evidence of ebolavirus infection in bats, China. Virol. J. 9, 236. doi:10.1186/1743-422X-9-236.

Zarocostas, J. (2020). Hope for the Ebola outbreak in DR Congo. Lancet 395, 773. doi:10.1016/S0140-6736(20)30495-5.

Zhou, P., Tachedjian, M., Wynne, J. W., Boyd, V., Cui, J., Smith, I., et al. (2016). Contraction of the type I IFN locus and unusual constitutive expression of IFN-α in bats. Proc. Natl. Acad. Sci. U.S.A. 113, 2696– 2701. doi:10.1073/pnas.1518240113.

