## Supplementary_figures_tables for "Marburg and Ebola virus infections elicit a complex, muted inflammatory state in bats"

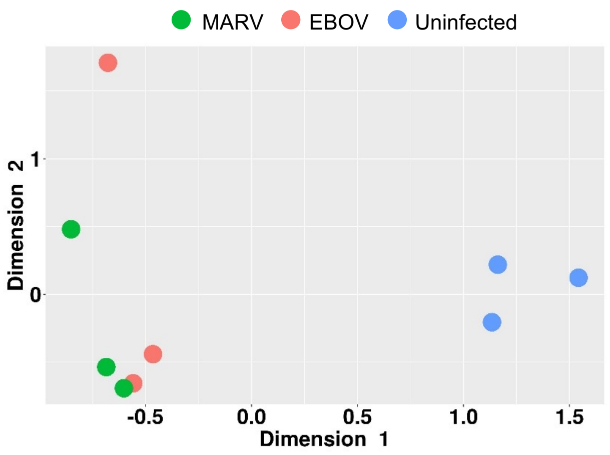


**Figure S1. A multidimensional scaling (MDS) plot of the merged gene expression data from kidneys of MARV-infected, EBOV-infected and uninfected bats.** The plot shows a clear separation between MARV infections, EBOV infections and the uninfected samples. The virus-specific signatures in this plot, along with that for livers and spleens (**Fig. 2**), demonstrate that the identified transcriptional responses to filovirus infections extends to the whole animal.

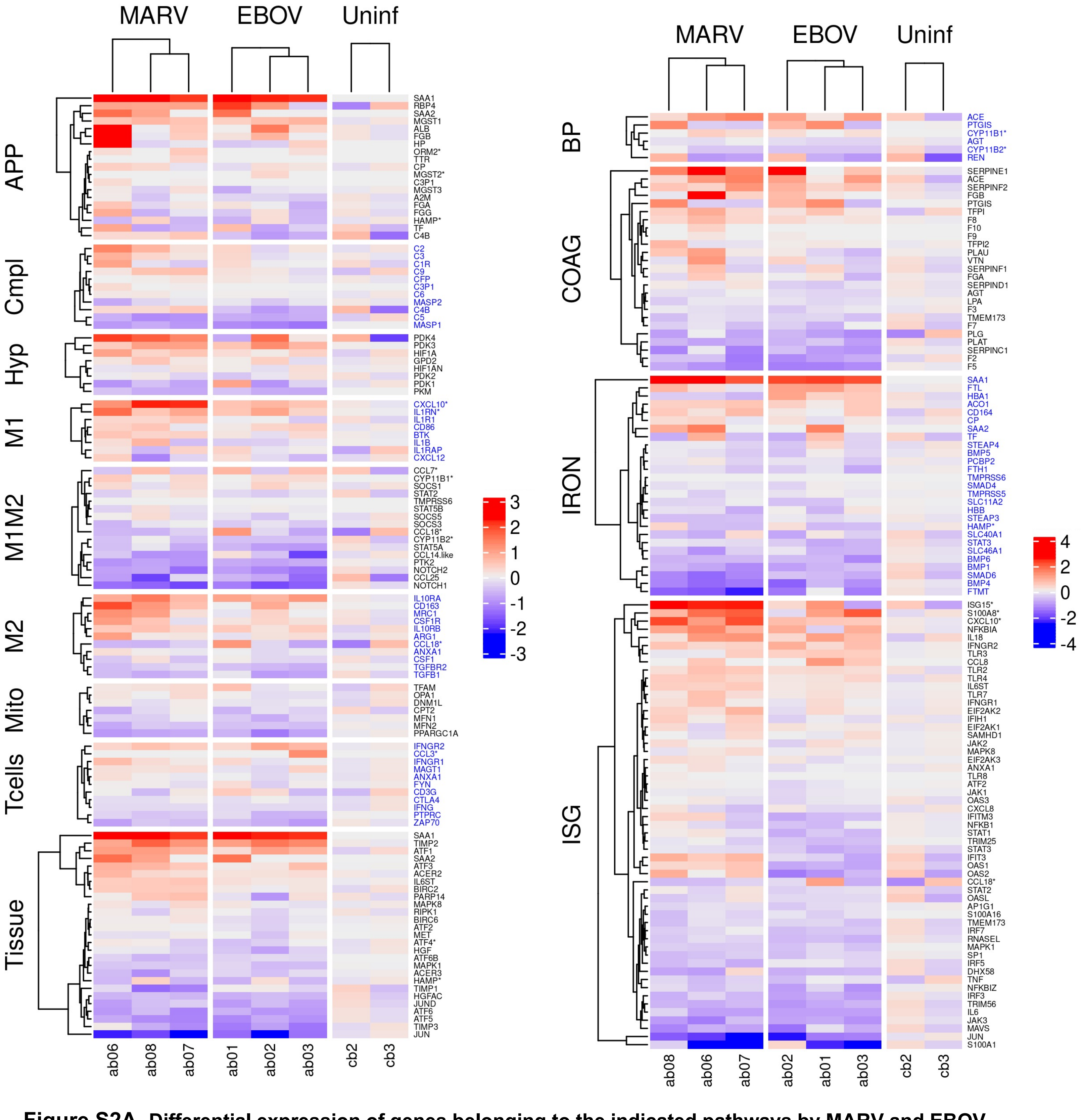


**Figure S2A. Differential expression of genes belonging to the indicated pathways by MARV and EBOV infections of bats in kidneys. Left panel:** genes related to acute phase response (APP), complement system (Cmpl), hypoxia (Hyp), tissue regeneration and apoptosis (Tissue), and genes specific to macrophages in the M1 state (M1), M2 state (M2) and, common to M1 and M2 states (M1M2). **Right panel:** genes related blood pressure (BP), coagulation (COAG), iron homeostasis (IRON) and interferon stimulated genes (ISG). The columns show samples from three MARV-infected bats, three EBOV-infected bats and two uninfected bats. The values are $\log_2$ of the fpm values, with the mean value of the samples from uninfected bats subtracted. The data show virus-specific responses which are not as pronounced as in the liver (**Fig. 4**); the responses are greater for MARV. Renin, in the BP group of gene, is only expressed in kidneys. There is a strong response in the SAA1/2 APP genes. ***** Genes whose bat versions are diverged from their human counterparts. Alternate blocks of gene names are colored black/blue to enable easy visual distinction of the blocks.

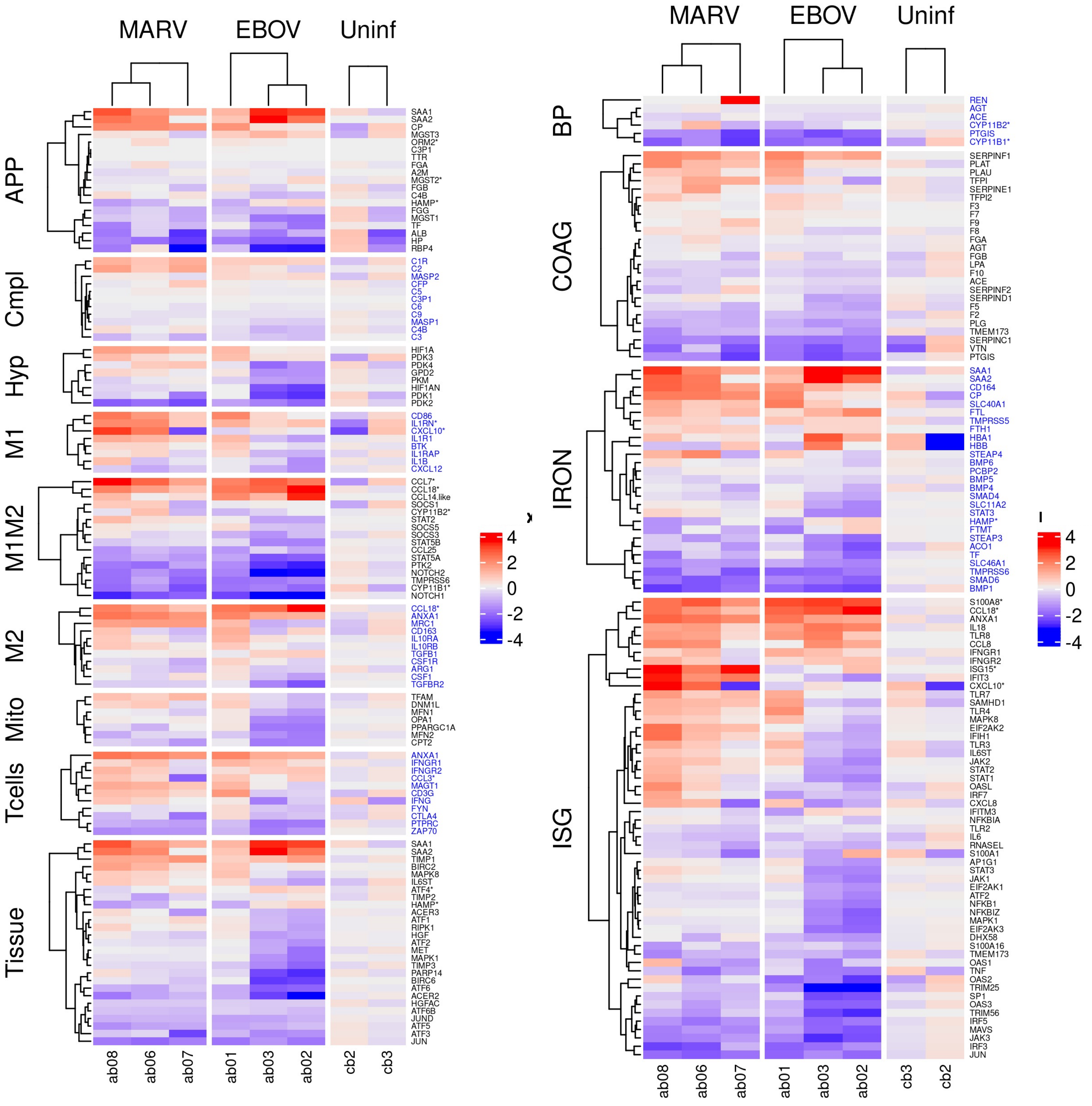


**Figure S2B. Differential expression of genes belonging to the indicated pathways by MARV and EBOV infections of bats in spleens. Left panel:** genes related to acute phase response (APP), complement (Cmpl) , hypoxia (Hyp), tissue regeneration / apoptosis (Tissue), and genes specific to macrophages in the M1 state (M1), M2 state (M2) and, common to M1 and M2 states (M1M2). **Right panel:** genes for blood pressure (BP), coagulation (COAG), iron homeostasis (IRON) and Interferon stimulated genes (ISG). The columns show the samples from three MARV-infected bats, three EBOV-infected bats and two uninfected bats. The values are $\log_2$ of the fpm values, with the mean value of the uninfected samples subtracted. The virus-specific response was not as pronounced as in the liver (**Fig. 4**) or kidneys (**Fig. S2A**), with larger effects in the case of MARV. A strong response in expression of the SAA1/2 APP genes was observed. Sting (TMEM173) expression was only detected in the spleen. * Genes whose bat versions are diverged from their human counterparts. Alternate blocks of gene names are colored black/blue to enable easy visual distinction of the blocks.

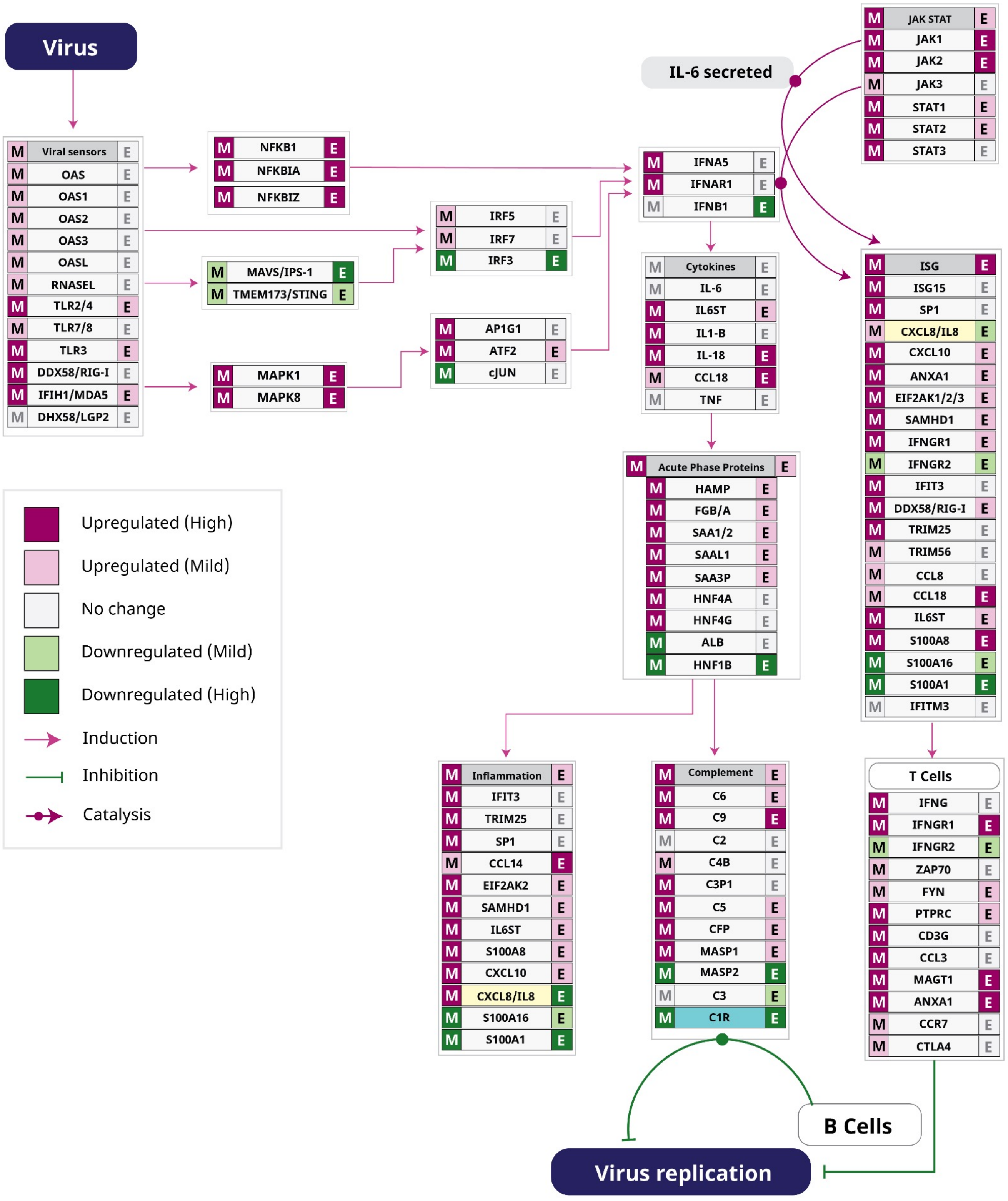


**Figure S3. Pathway analysis of innate response to filovirus infections in bats.** Viral RNA and proteins are detected by innate immune sensors, leading to upregulation of type I interferons (IFN$\alpha$ and IFN$\beta$) and interferon stimulated genes (ISG) through the JAK-STAT pathway. Cytokines and chemokines create an anti-viral state in the cell. Inflammation (in part mediated by IL-6) triggers expression of acute phase proteins (APPs). Secretion of interferon-**γ** gamma enables an adaptive immune response. In human and bat cells interferon responses to the filoviruses were mostly similar, with a few virus-specific differences. VP35 from both viruses interferes with IRF3/7, while EBOV VP24 inhibits STAT1, and MARV-VP40 inhibits JAK1, which can lead to differences in responses to MARV and EBOV. The robust innate response to filovirus infection in bats and humans suggests any constitutive expression of innate response genes in bats is unlikely to be relevant to explain the difference in the pathogenesis. **Fig. 4** shows the relative changes in expression upon infection for these genes. Here, colored bands flanking gene names depict the effect of filovirus infection on gene expression for MARV (left) and EBOV (right).

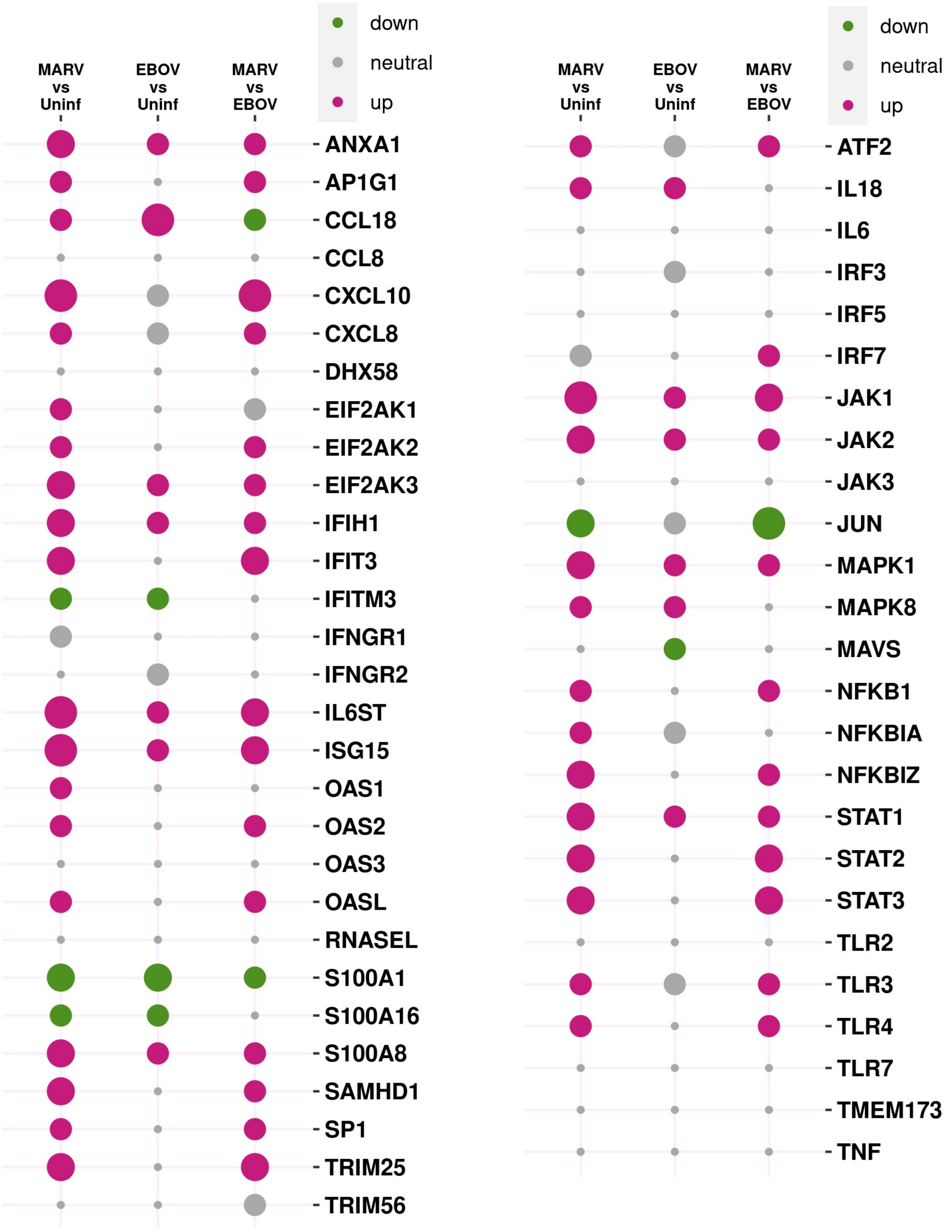


**Figure S4. Induction of interferon stimulated genes (ISG) in livers of filovirus-infected bats.** MARV and EBOV elicit a strong innate response driven by IFNs. The balloon plot shows comparisons of responses of ISG genes to MARV and EBOV against uninfected animals and against each other. The radius of circle is proportional to $\log_2$(ratio), red is $\log_2$(ratio) > 0.6, green is $\log_2$(ratio) < -0.6, gray is $-0.6<\log_2$(ratio)$<0.6$. The response is stronger in the MARV infected animals, corresponding to the greater replication of MARV as compared to EBOV in ERBs.

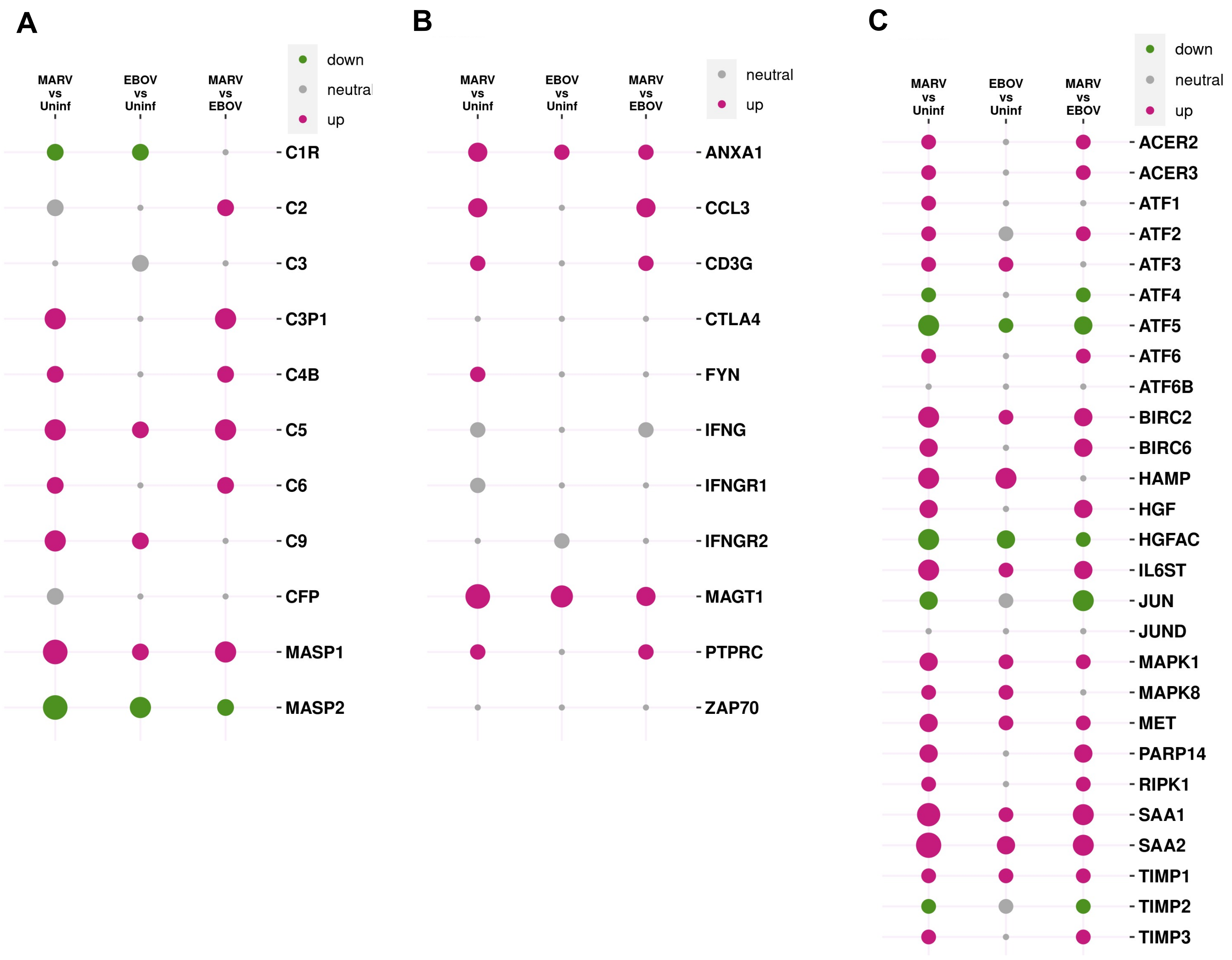


**Figure S5. Tissue regeneration, the complement system and CD8+ T cells.** Responses of genes in MARV and EBOV infected bats against uninfected bats and against each other. Balloon plots with the radius of circles proportional to $\log_2$ (ratio), gray is used when absolute values of $\log_2$ (ratio) < 0.6. **A.** The complement system: filovirus infections downregulated C1R, C3 and MASP2, likely leading to reduced antibody activities. **B.** CD8+ T cell genes: most of them were upregulated by filovirus infections, indicating at CD8+ T cell response, which is likely to be involved in clearance of filovirus infections in bats **C.** Tissue regeneration: filovirus infections in bats eventually trigger tissue regeneration, reflected in the activation of M2 macrophages, which are anti-inflammatory and promote regeneration of tissues. The greater effect of MARV infection as compared to EBOV is consistent with the higher viral loads..

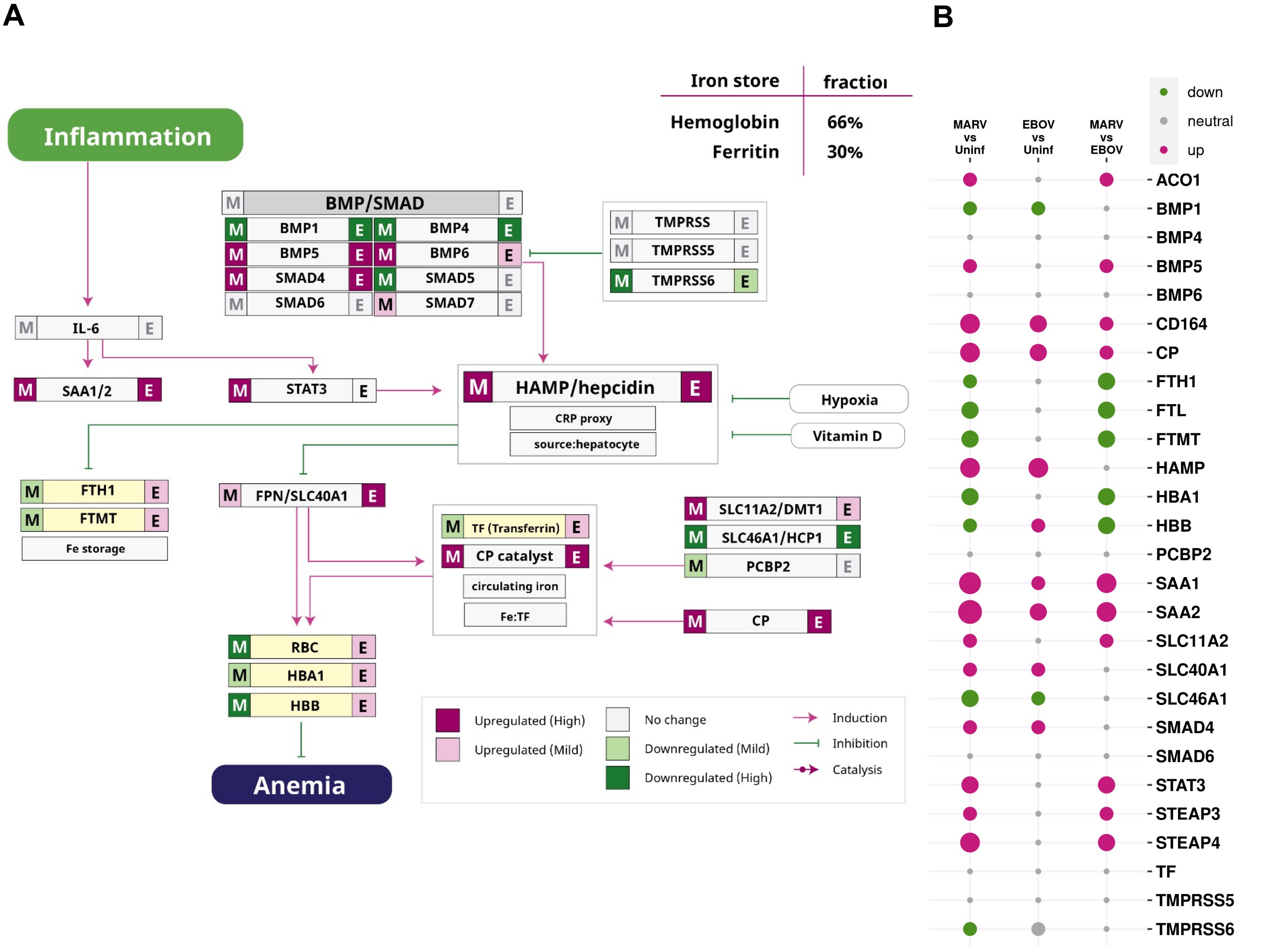


**Figure S6. Iron metabolism. A.** Pathway analysis of genes involved in iron homeostasis. Colored bands flanking gene names depict the effect of filovirus infection on gene expression for MARV (left) and EBOV (right), with upregulation depicted by shades of red and downregulation depicted by shades of green. The lines show induction (red) and inhibition (green). Filovirus infections lead to upregulation of HAMP. In MARV infection, this may lead to a decrease in blood iron levels and impaired hematopoiesis, but in EBOV infection, this regulation seems to be broken, leading to a high iron state. These differences between MARV and EBOV infections are consistent with the greater replication of MARV in ERBs. ACO1 is upregulated during filovirus infections, suggesting the cytosol has abundant iron. **B.** The balloon plot compares responses of genes in MARV and EBOV infected bats against uninfected bats and against each other. The radius of circle is proportional to $\log_2$ (ratio), gray is used when absolute values of $\log_2$(ratio) < 0.6.

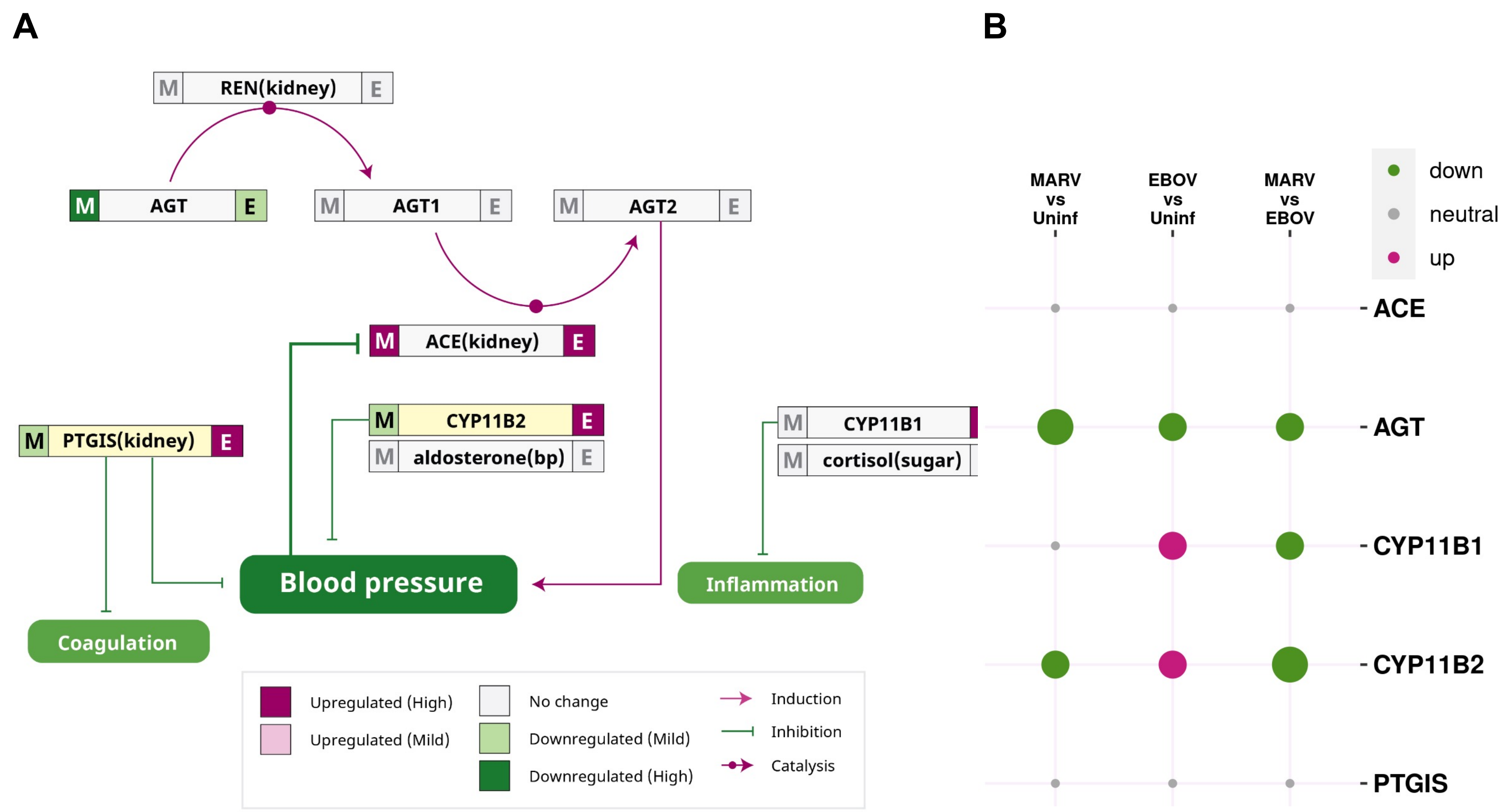


**Figure S7. Blood pressure pathways**. **A.** Analysis of pathways associated with renin (REN) and angiotensin I-converting enzyme (ACE) which catalyze conversion of AGT to AGT2, which in turn constricts blood vessels to increase pressure. Filovirus infections in ARBs lowered expression of AGT but increased PTGIS, which reduces blood pressure and coagulation and increased ACE, which is a sensor of low blood pressure. Thus, filovirus-infected bats induces an anti-coagulative state with low blood pressure. The colored bands on either side of the gene names depict the effects of MARV (left) and EBOV (right) infections on expression of the indicated genes. **B.** The balloon plot compares responses of genes to MARV and EBOV compared to uninfected animals and against each other. The radius of circle is proportional to $\log_2$ (ratio), gray is used when absolute values of $\log_2$ (ratio) < 0.6.

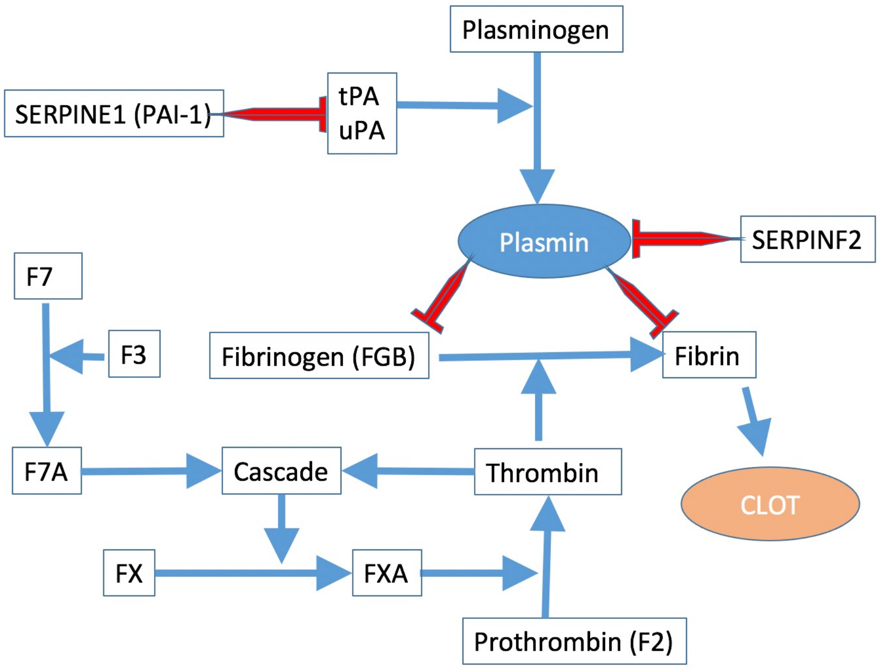


**Figure S8. The coagulation pathway.** The coagulation pathway involves a cascade of activations that eventually results in fibrin formation that is part of the clot. There is an opposing process where plasmin degrades fibrinogen and fibrin thereby dissolving clots. The figure only shows processes that are relevant to the discussion in the paper. The red lines show inhibition of a process or degradation, while the blue arrows signify enhancement of the process/product. The process starts with tissue factor (F3) activating F7 (to form F7A) which eventually leads to the formation of Thrombin, which facilitates fibrin synthesis**. Fig. S9** shows the expression changes in coagulation pathway genes upon filovirus infections.

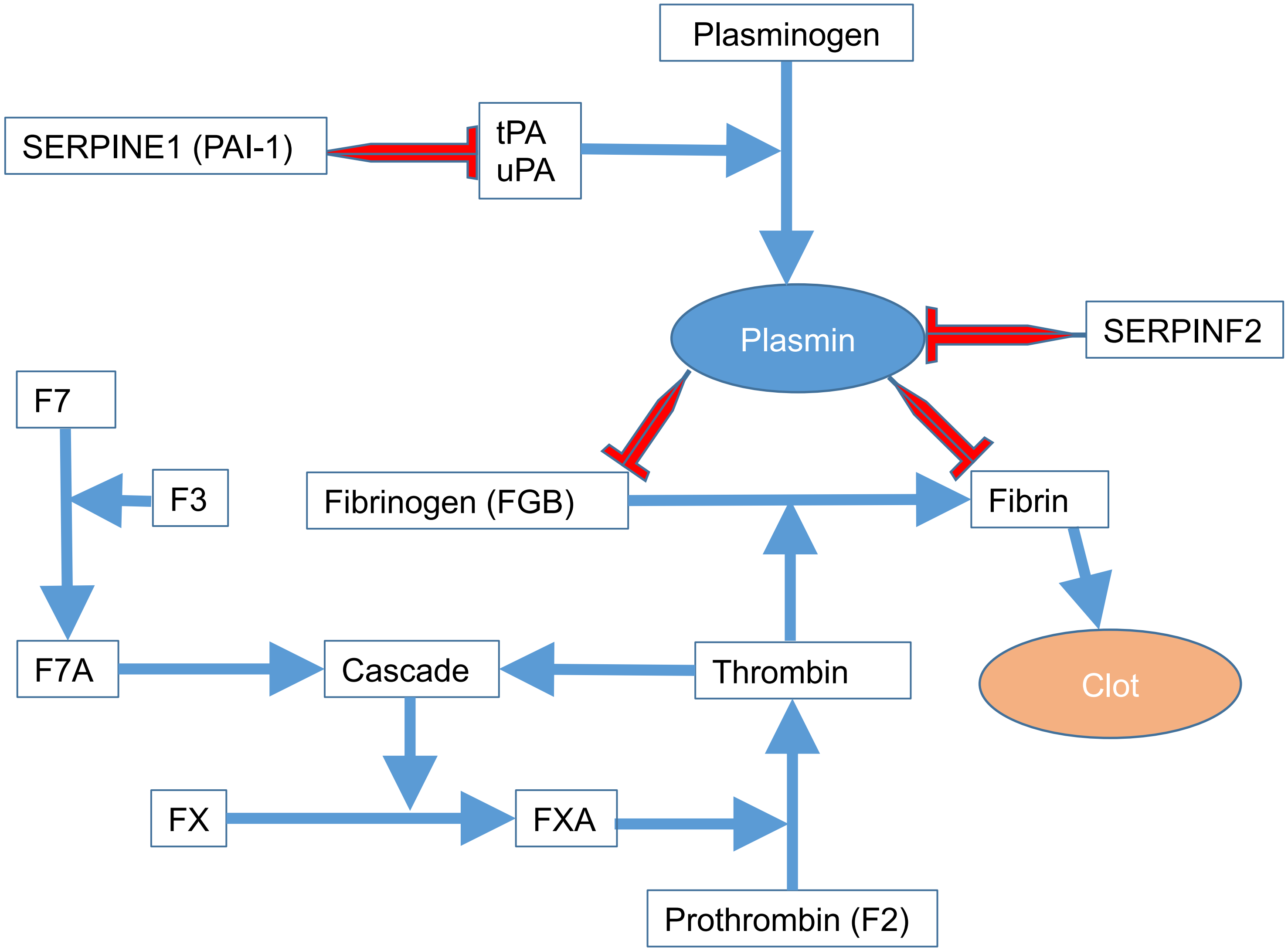


**Figure S8. The coagulation pathway.** The coagulation pathway involves a cascade of activations that eventually results in fibrin formation that is part of the clot. There is an opposing process where plasmin degrades fibrinogen and fibrin thereby dissolving clots. The figure only shows processes that are relevant to the discussion in the paper. The red lines show inhibition of a process or degradation, while the blue arrows signify enhancement of the process/product. The process starts with tissue factor (F3) activating F7 (to form F7A) which eventually leads to the formation of Thrombin, which facilitates fibrin synthesis**. Fig. S11** shows the expression changes in coagulation pathway genes upon filovirus infections.

A

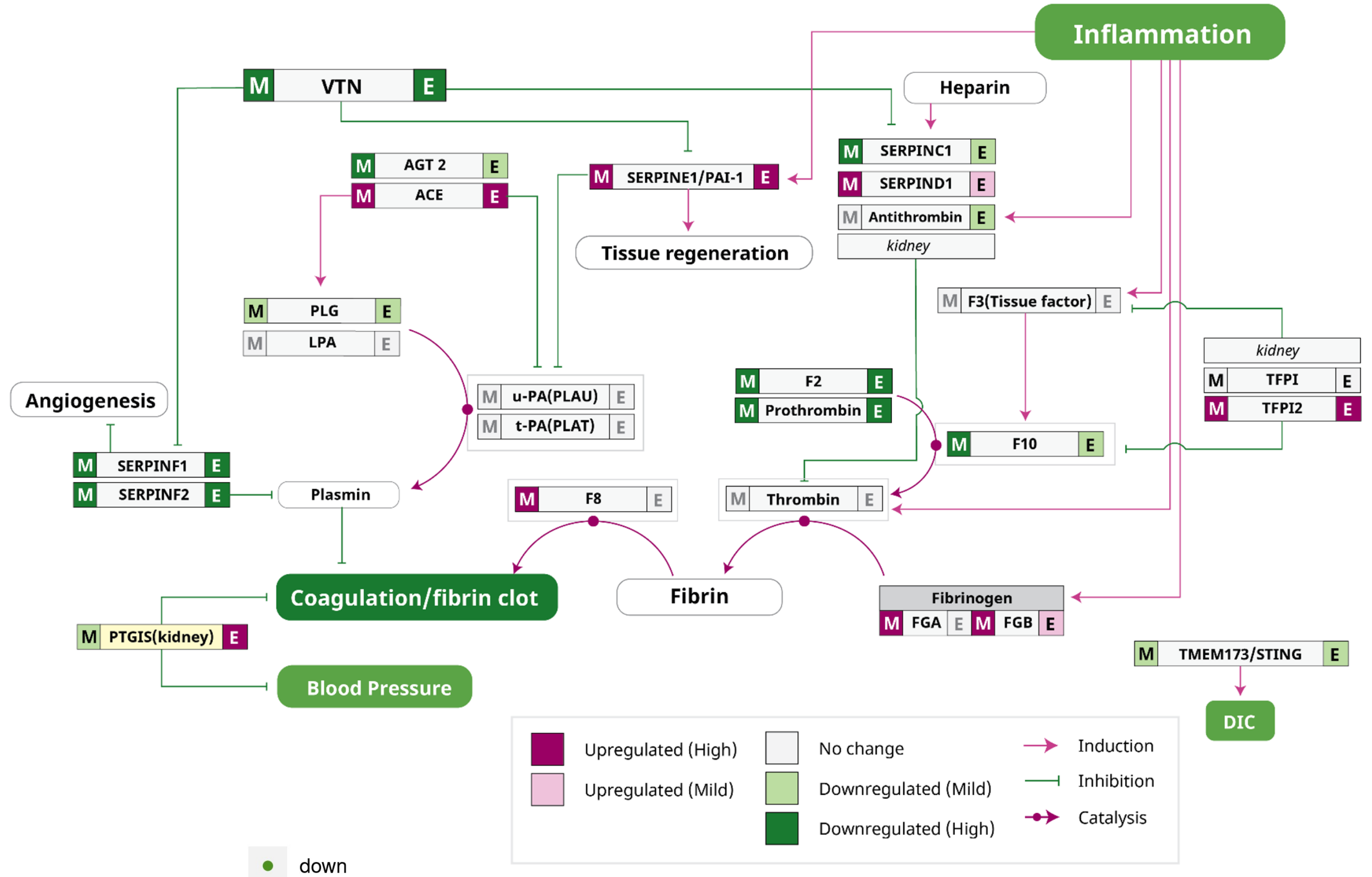


B

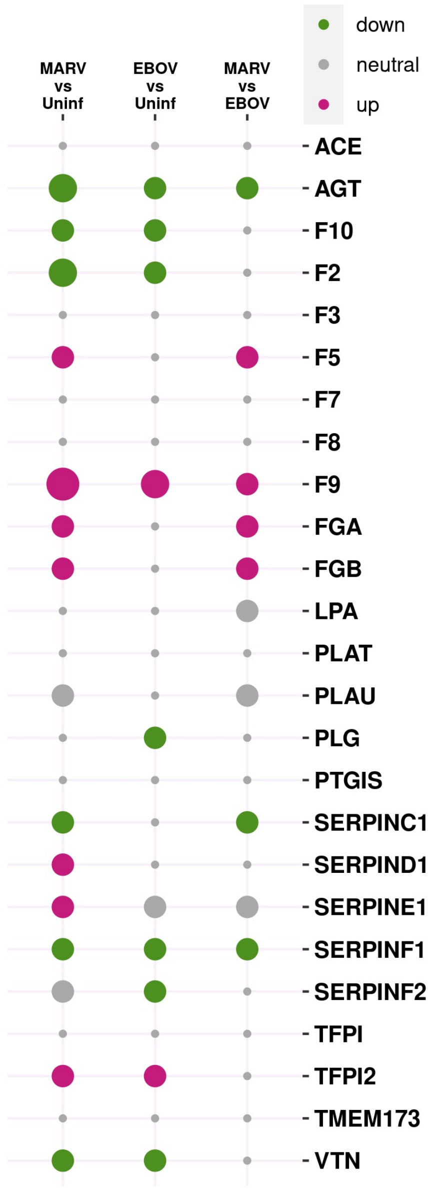


**Figure S9. Coagulation pathway. A.** Analysis of pathways associated with coagulation cascade (shown in **Fig. S8**). Except for F3, the tissue factor, all other proteases in the cascade require activation. The serine protease plasmin opposes this process by attacking the fibrin mesh to dissolve clots. Urokinase-type plasminogen activator (uPA, PLAU gene) activates plasminogen to generate plasmin. SERPINE1 (PAI-1) inhibits the activity of uPA, blocking the creation of plasmin, thereby stabilizing clots. SERPINF2 also stabilizes clots by directly inhibiting plasmin. PTGIS creates prostacyclin, which prevents platelet aggregation, yet another path for inhibiting clot formation and coagulation. Prostacyclin is also a potent vasodilator. Failure in the control of plasmin (e.g., inactivation of SERPINF2 which inhibits plasmin) can lead to hemorrhagic diathesis, while blocking plasmin activity can lead to excessive clotting. Filovirus infections in bats lead to low F2 (which lowers fibrin) and raise levels of PTGIS, which suggest that filovirus-infected bats are in a low coagulation state. **B.** The balloon plot compares responses of genes to MARV and EBOV compared to uninfected animals and against each other. The colored bands on either side of the gene names depict the effects of filovirus infections on gene expression for MARV (left) and EBOV (right) compared to uninfected animals. The radius of circle is proportional to $\log_2$ (ratio), gray is used when absolute values of $\log_2$ (ratio) < 0.6.

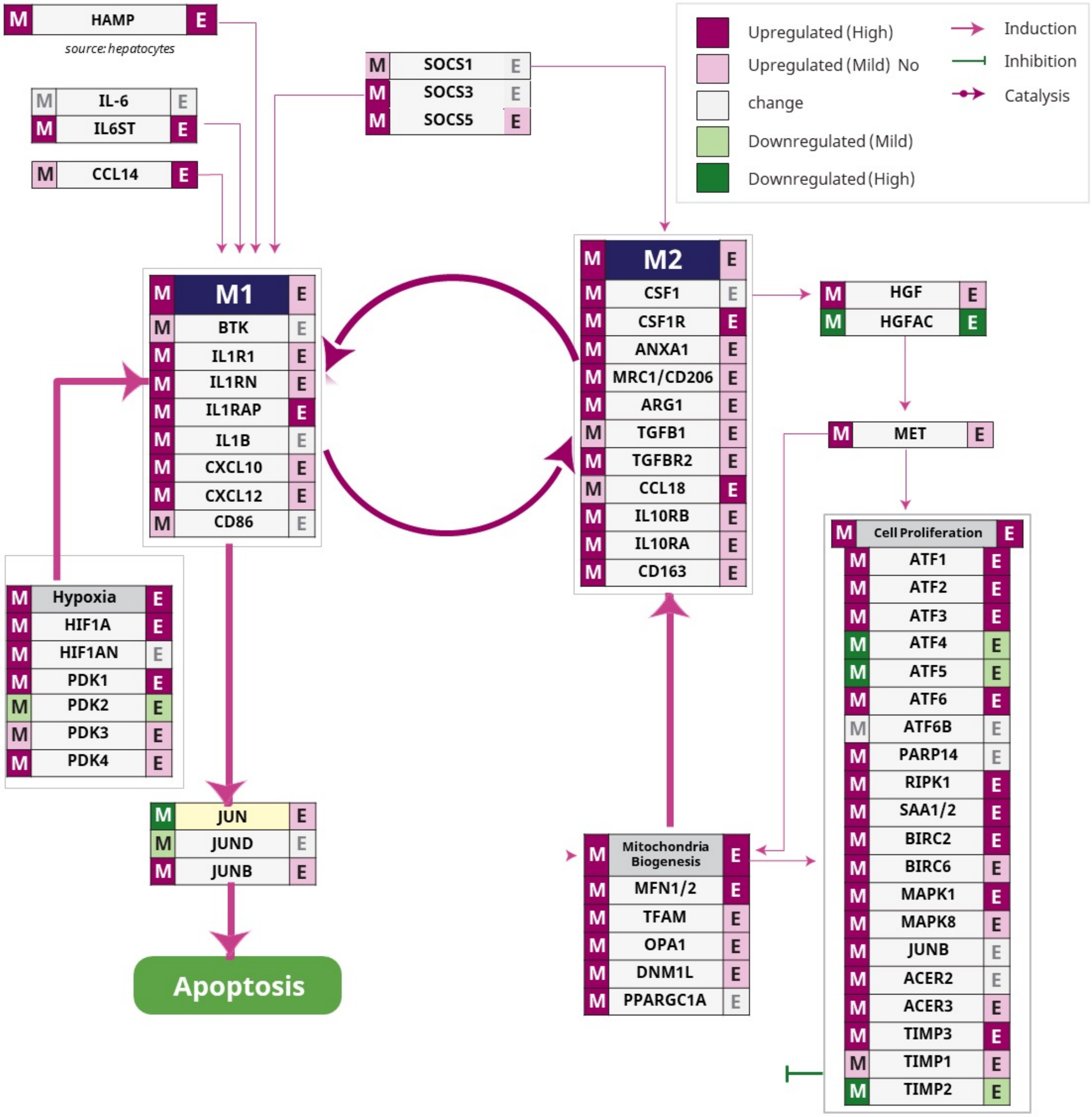


**Figure S10. Pathway analysis related to macrophage polarization during filovirus infections.** Differential expression of genes involved in macrophage polarization between the pro-inflammatory M1 state and the anti-inflammatory M2 state. Colored bands flanking gene names depict the effects of MARV (left) and EBOV (right), with upregulation depicted by shades of red and downregulation depicted by shades of green. The lines show induction (red) and inhibition (green). Filoviral infections initially lead to proinflammatory state of macrophages (M1) which then transitions to anti-inflammatory state (M2). The differential expression levels of these genes as balloon plots is shown in Fig. S5. HAMP induces macrophages in the proinflammatory M1 state which phagocytose infected cells and induces apoptosis. In contrast, the M2 macrophages induce cell proliferation and tissue regeneration. Different SOCS family molecules serve as molecular switches that control M1/M2 macrophage switch. Fatty acid oxidation and increased mitochondrial numbers and activity indicates an M1 to M2 switch. During filoviral infections, both M1 and M2 states are seen, based on the markers specific to them, but the ratio of M1 to M2 was skewed more towards M2 in EBOV-infected bats case, consistent with the more limited virus replication, while the M1 to M2 transition was still underway in the MARV-infected bats. The switch to the M2 state is probably the key to the resilience of bats during filovirus infections, allowing bats to tolerate filovirus infections without significant adverse effects. There is also a connection between iron metabolism and macrophage M1/M2 polarization with increasing iron (Fig. S6) favoring the M2 state.

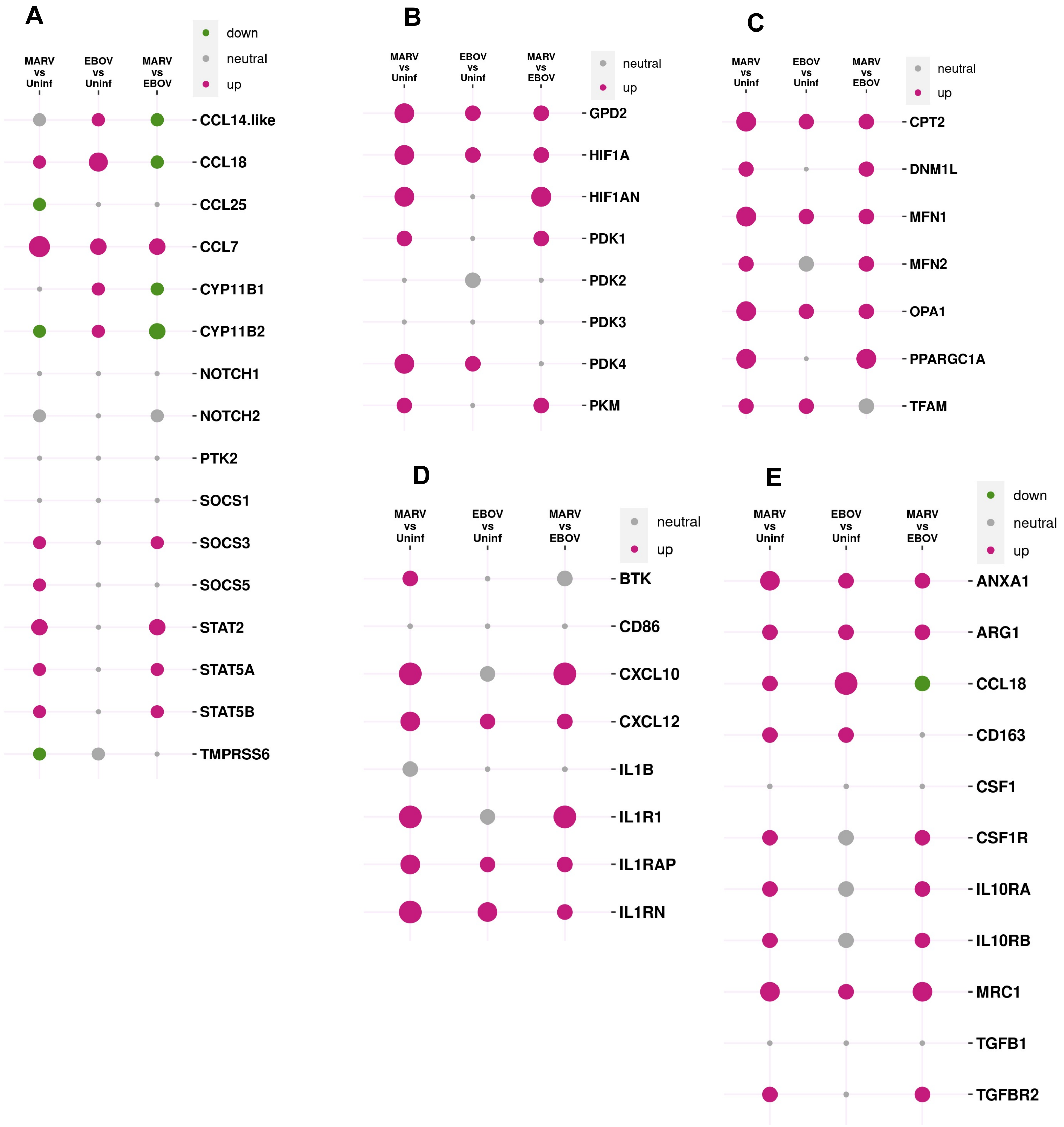


**Figure S11. Macrophage activation.** Left panel(**A**): genes common to both M1 and M2 states. Top right panels (**B,C**): genes involved in hypoxia and mitochondrial respiration/abundance. Bottom right panels(**D,E**): genes specific to the M1 and M2 macrophages which show that both M1 and M2 states are activated by filovirus infections with the M1/M2 ratio being higher in MARV infection. The balloon plots shows comparisons of gene expression responses to MARV and EBOV in comparison with uninfected animals. The radius of circle is proportional to $\log_2$ (ratio), gray is used when absolute values of $\log_2$ (ratio) < 0.6. There are more activated macrophages in the MARV-infected bats compared to EBOV-infected bats, consistent with greater level of MARV replication.

**Table ST1. List of bat samples profiled using mRNA-seq**

| Tissue | Uninfected | MARV | EBOV |
|---|---|---|---|
| Liver | 3 | 3 | 3 |
| Lung | 0 | 3 | 3 |
| Spleen | 3 | 3 | 3 |
| Kidney | 3 | 3 | 3 |
| Large intestine | 0 | 3 | 3 |

**Table ST2. MARV transcripts detected using mRNAseq in tissues of MARV-infected bats**

| Organ | ab07 | ab08 | ab06 |
|---|---|---|---|
| **Liver** | 79 | 8.7 | 3.1 |
| Spleen | 56 | 8.5 | 3.4 |
| Large intestine | 2.3 | 10.1 | NA |
| Lung | 2.3 | 0 | NA |

NA, not analyzed, stands for samples that failed mRNAseq. The values are expressed in tpm. EBOV transcripts are seen only in liver in very low numbers.

**Table ST3A. Divergent pathways and genes upregulated by MARV and EBOV infection: vascular, mitochondrial, oxidation-reduction and innate immunity**

| Gene | Process |
|---|---|
| | Vascular |
| HAMP/ Hepcidin | Cellular iron ion homeostasis inflammatory induces M1 macrophages |
| SRGN | Platelet degranulation |
| AK3 | Platelet production nucleoside diphosphate phosphorylation mitochondrial |
| GLRX | Antioxidant defense system VEGF expression vascular growth |
| CEBPZOS | Blood cell maturation |
| | Mitochondrial |
| MRPL50 | Organelle organization |
| GSTZ1 | Detox reduces oxidative stress |
| | Redox |
| PECR | Oxidation-reduction process lipid synthesis regeneration cell growth |
| GALM | Glucose metabolic process |
| | Macrophages |
| APOL6 | Monocyte to macrophage differentiation lipid metabolism |
| TIMD4 | Expressed by macrophage maintains killer T cell activity |
| CCL18 | Attracts T cells to macrophages cellular response to IFNG |
| CXCL10 | secreted by macrophage in response to IFNG role in hypertension Immune response |
| CCL3 | Macrophage inflammatory protein cellular response to IFNG |
| PRXL2A | Inhibits production of inflammatory cytokines by macrophages |
| S100A12 | paralog of S100A8 IL-10 induced monocytes macrophages Proinflammatory mast cell chemoattractant innate immune response |
| PLAC8 | Expressed by macrophage phospholipid metabolic process |
| ISG15 | Regulation of IFNG production Mito function in macrophages |
| CCL7 | Regulates macrophages attracts monocytes eosinophils but not neutrophils cellular response to interferon-gamma |
| | Innate immunity |
| CLEC4F | Endocytosis pathogen detector |
| CLECL1 | Expressed by dendritic and B cells enhances IL-4 production regulates immune response |
| CXCL3 | Chemoattractant for neutrophils immune response |

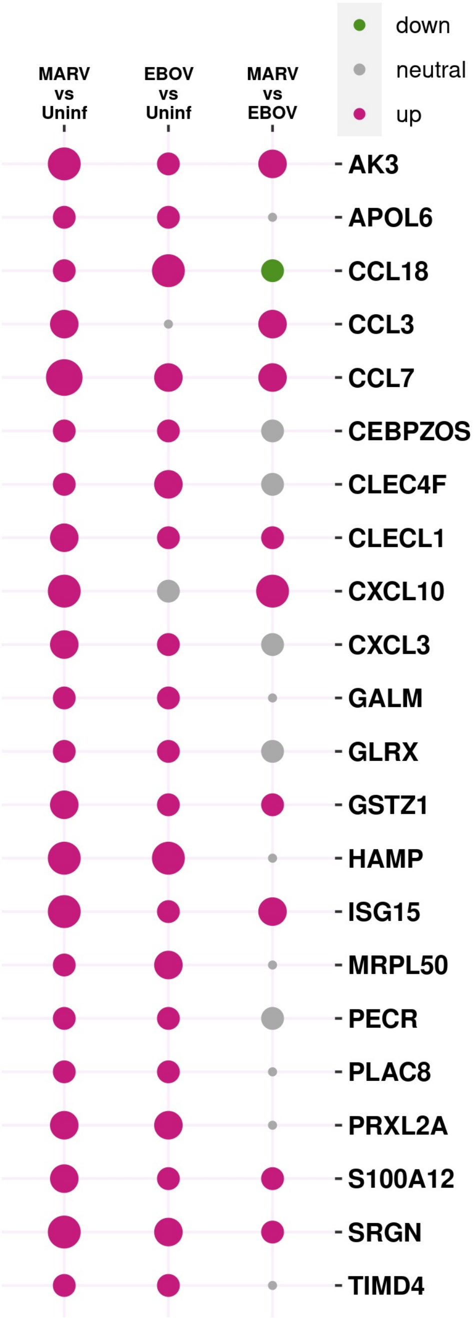


The balloon plot on the right compares responses of genes to MARV-, EBOV- infected and uninfected samples against each other. The radius of circle is proportional to $\log_2$ (ratio), red is for positive, green is for negative values and gray is used when absolute values of $\log_2$(ratio) < 0.6.

### Table ST3B. Divergent pathways and genes upregulated by MARV and EBOV infection: T cell activity, complement, digestion, toxins, inflammation and tissue regeneration

| Gene | Process |
|---|---|
| | T cell |
| LY6E | T cell development negative regulator of monocytes dampens response |
| | Complement |
| MBL2 | Activates complement mannose-binding innate immune response |
| | Digestion |
| PRB1 | Digestion salivary proline-rich protein |
| | Toxins |
| UGT2A3 | Excretion of toxic compounds flavonoid biosynthetic process |
| | Inflammation |
| ORM2 | Acute phase reactant regulation of immune system process |
| IL1RN | IL-1R antagonist inhibits IL1A/B, modulates inflammation immune response |
| MGST2 | Generates LTC4 mediates stress |
| S100A8 | Leukocyte migration involved in inflammatory response |
| | Tissue regeneration |
| ACOT13 | Essential for cell proliferation mitochondrial function (macrophage M2) |
| S100P | Cell proliferation, response to organic substance |
| NUDT16 | Positive regulation of cell proliferation |
| EIF4EBP3 | Negative regulation of translational initiation |
| | Other |
| ZG16B | Atherosclerosis retina homeostasis |
| SUB1 | Regulation of transcription from RNA polymerase II promoter |
| UNDEF108 | |
| UNDEF771 | |

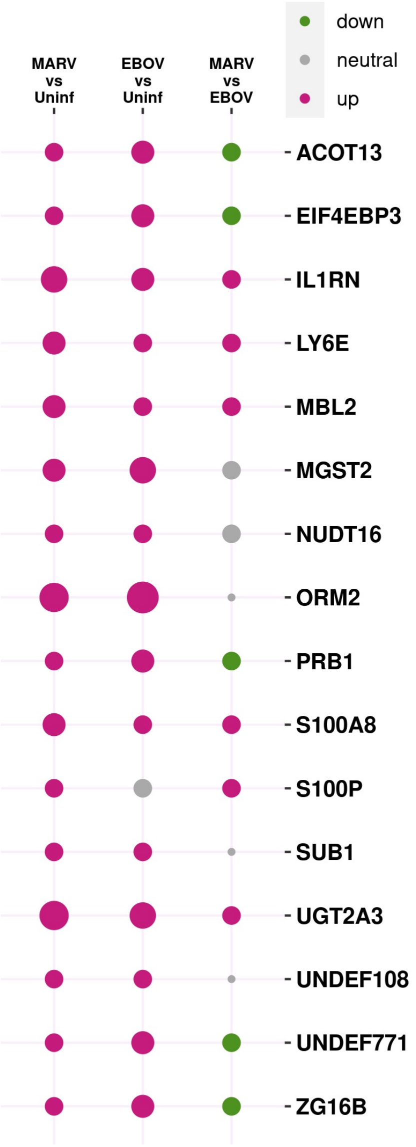


The balloon plot at the right compares responses of genes to MARV-, EBOV- infected and uninfected samples against each other. The radius of circle is proportional to $\log_2$ (ratio), red is for positive, green is for negative values and gray is used when absolute values of $\log_2$(ratio) < 0.6.

**Table ST4A. Divergent pathways and genes downregulated by MARV and EBOV infection: mitochondrial activity, vascular function, inflammation, innate immunity, lipids, toxins and macrophage activation**

| Gene | Process |
|---|---|
| | Mitochondria/oxidation/fatty-acid |
| MRPL54 | Organelle organization |
| CA3 | Nitrogen metabolism bicarbonate transport |
| RNASEH1 | mtDNA replication mutations lead to autoimmunity(T1D) |
| CHCHD7 | Protein import |
| CHCHD10 | Negative regulation of ATP citrate synthase activity |
| | Vascular |
| ART4 | Blood group antigens protein ADP-ribosylation |
| HRG | Platelet degranulation Histidine-rich glycoprotein |
| MEG3 | Negative regulation of VEGF receptor signaling pathway |
| TMEM80 | Increased transferrin (TF) endocytosis |
| | Inflammation |
| AHSG | Regulation of inflammatory response |
| CRELD2 | ER-stress response |
| TEX264 | Stress , elevated platelet cytosolic Ca2+ responsive |
| | Innate immunity |
| BTBD6 | Class I MHC-mediated antigen processing/presentation |
| C8G | Complement Complement component C8 gamma chain |
| EEF1D | Positive regulation of I-kB kinase/NF-kB signaling |
| MIIP | Down-regulates NFKB2 and ICAM1 inhibition of migration/invasion |
| TMEM80 | Increased vaccinia virus (VACV) infection Decreased NF-kB reporter expression |
| CRIP1 | Cysteine-rich protein 1 |
| | Lipids |
| MOGAT1 | Triacylglycerol biosynthesis and Metabolism |
| | Toxin |
| GSTA5 | Detoxification glutathione metabolic process |
| SLC17A2 | Sodium/anion cotransporter family ossification |
| | Macrophages |
| RNASET2 | Chemoattractants for macrophages and modulate the inflammatory processes |
| | Vascular (iron) |
| ATP6V0D2 | Cellular iron ion homeostasis |
| BSG | Carries OK antigens on red blood cells:cell surface receptor Signaling pathway:inflammation |
| LCN2 | Sequesters iron (antibacterial) iron/toxin transport cisplatin resistance innate immune response |

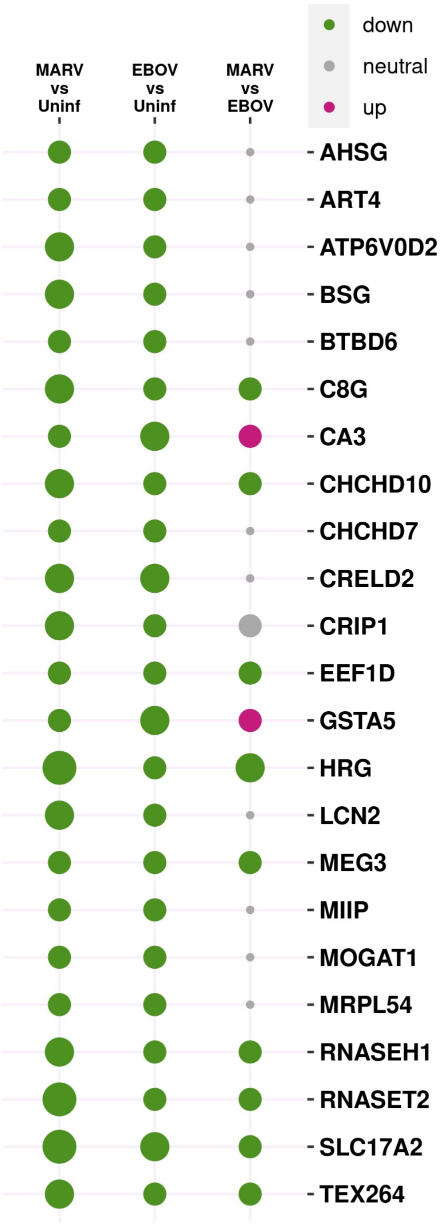


The balloon plot at the right compares responses of genes to MARV-, EBOV- infected and uninfected samples against each other. The radius of circle is proportional to $\log_2$ (ratio), red is for positive, green is for negative values and gray is used when absolute values of $\log_2$(ratio) < 0.6.

**Table ST4B. Divergent pathways and genes downregulated by MARV and EBOV infection: splicing, T cell activity, metabolism, digestion, macrophages and apoptosis**

| Gene | Process |
|---|---|
| | Splicing |
| SNRPN | mRNA splicing via spliceosome |
| | T cell |
| GPX4 | Role in primary T-cell response to viral infection protects T-cells from ferroptosis supports T-cell expansion mitochondrial |
| | Metabolism |
| TSTD1 | Thiosulfate sulfur transferase/rhodanese-like |
| NUDT14 | Hydrolyzes UDP-glucose to glucose 1-phosphate and UMP and ADP-ribose to ribose 5-phosphate and AMP |
| | Digestion |
| FABP2 | Fatty acid-binding protein intestinal |
| | Macrophages |
| APOC1 | Activated when monocytes differentiate to macrophages<br><br>positive regulation of cholesterol esterification |
| | Apoptosis/tissue regeneration |
| MSTO2P | Proliferation |
| LCMT1 | Regulation of apoptotic process |
| PRDM11 | Inhibits proliferation induces apoptosis |
| | Other |
| SPP2 | Negative regulation of endopeptidase activity |
| SCGB1C1 | Upper respiratory tract |
| RAMP1 | Regulation of GPCR signaling pathway |
| S100G | Calcium ion/vitamin D binding mineral absorption |
| RANGRF | Positive regulation of GTPase activity |
| RPS28 | Viral process |
| BEX4 | |
| URAHP | |
| C19orf12 | |
| FLJ37453 | |
| TMEM141 | |
| FXYD1 | Positive regulation of sodium ion export |
| UNDEF425 | |
| UNDEF464 | |

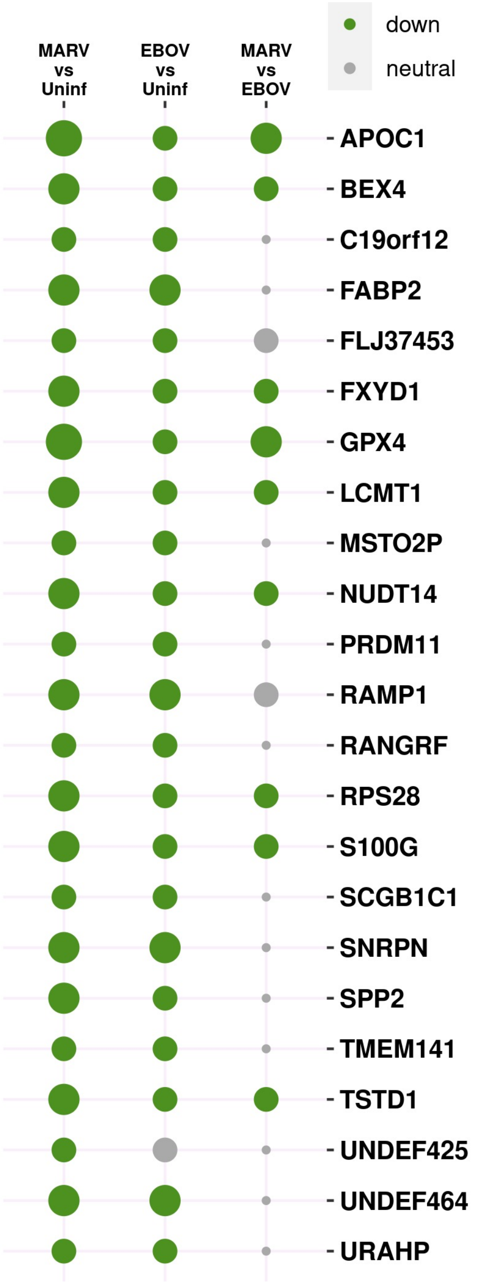


The balloon plot compares responses of genes to MARV-, EBOV- infected and uninfected samples against each other. The radius of circle is proportional to $\log_2$ (ratio), red is for positive, green is for negative values and gray is used when absolute values of $\log_2$ (ratio) < 0.6.

**Table ST5. Divergent pathways and genes upregulated by MARV but not EBOV infection: macrophages, complement, apoptosis, mitochondrial respiration, innate immunity, inflammation, digestion, T cells and the vascular system**

| Gene | Process |
|---|---|
| | Macrophages |
| BPI | Negative regulation of IL-6 production expressed by Macrophages Bactericidal permeability-increasing protein |
| | Complement |
| CD46 | Inactivates C3b and C4b protect host cell from damage by Complement innate immune response |
| | Apoptosis |
| XAF1 | Response to interferon-beta proapoptotic |
| TNFRSF10A | Activation of NF-kB-inducing kinase activity cell apoptosis |
| BID | Positive regulation of apoptotic process |
| | Mitochondria/glycolysis/fatty-acid |
| SPR | Oxidoreductase activity and aldo-keto reductase (NADP) activity Nitric oxide biosynthetic process |
| ECHDC3 | Fatty acid pathways( Macrophage M2) |
| | Innate immunity |
| IFI30 | Antigen processing IFNG-mediated signaling pathway |
| TRIM22 | IFNG-mediated signaling pathway antiviral ubiquitinates viral proteins |
| ICAM1 | IFNG-mediated signaling pathway |
| | Inflammation |
| IL33 | Positive regulation of inflammatory response |
| | Digestion |
| SULT2A1 | Digestion Bile salt sulfotransferase |
| | T cells |
| CMTM6 | Protects PD-L1 inhibits T cells |
| GZMH | Immune response T cell Granzyme H |
| | Vascular |
| EMP2 | Positively regulates VEGF-A , integrin-mediated signaling pathway |
| RHOG | Platelet activation Rho-related |
| | Other |
| ARF1 | Viral process ADP-ribosylation factor 1 |
| RBM12B-AS1 | RBM12B antisense RNA 1 |
| RNF213 | Protein ubiquitination E3 ubiquitin-protein ligase RNF213 |
| UNDEF25 | |
| CUNH17of100 | |
| UNDEF113 | |

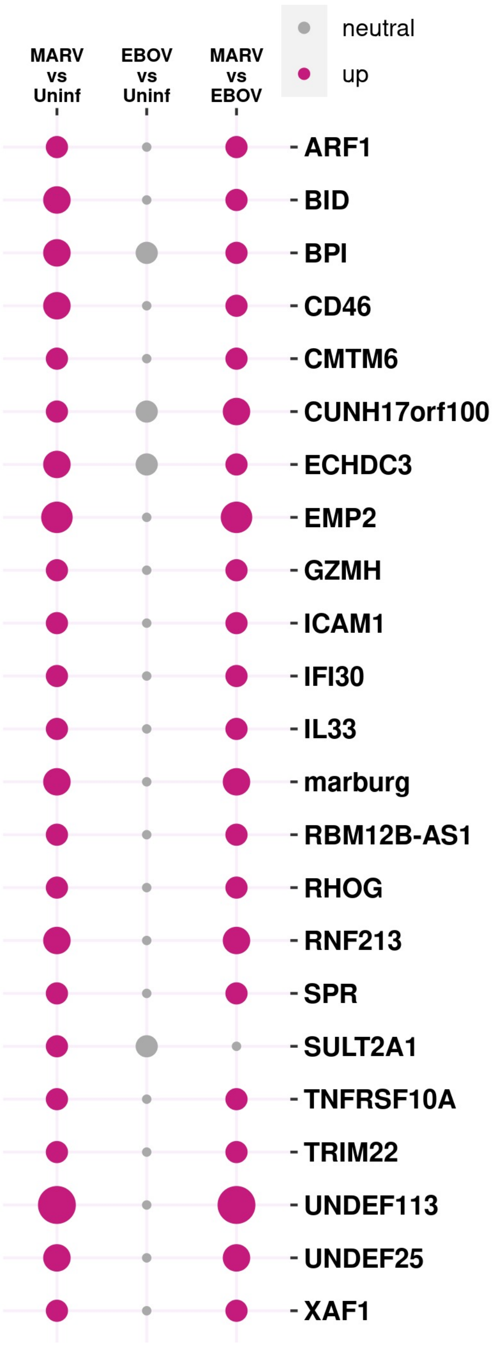


The balloon plot at the right compares responses of genes to MARV-, EBOV- infected and uninfected samples against each other. The radius of circle is proportional to $\log_2$ (ratio), red is for positive, green is for negative values and gray is used when absolute values of $\log_2$ (ratio) < 0.6.

**Table ST6. Divergent pathways and genes downregulated by MARV but not EBOV infection: mitochondrial, vascular, inflammation, digestion, innate immunity, complement, apoptosis and splicing**

| Gene | Process |
|---|---|
| | Mitochondrial |
| MRPL41 | Translation |
| NDUFV3 | Mitochondrial electron transport NADH to ubiquinone |
| MDH2 | Gluconeogenesis mitochondrial |
| TOMM6 | Protein targeting to mitochondrion |
| NDUFB11 | Respiratory electron transport chain |
| ATP5MPL | Mitochondrial membrane atp synthase |
| NDUFB1 | Mitochondrial electron transport NADH to ubiquinone |
| NDUFA4 | Mitochondrial electron transport NADH to ubiquinone |
| COA3 | Positive regulation of mitochondrial translation |
| UQCRBP1 | Ubiquinol-cytochrome c reductase binding protein pseudogene 1 |
| | Inflammation/oxidative stress |
| NDUFB4 | Response to oxidative stress |
| | Digestion |
| AMY2B | Digestion Alpha-amylase 2B |
| | Vascular |
| SERPINA4 | Negative regulation of endopeptidase activity |
| ZNF24 | Represses VEGF vascularization myelination |
| | Innate immune(complement) |
| C4BPB | Innate immune response |
| CCL16 | Cellular response to interferon-gamma |
| RPS27A | Toll-like receptor signaling pathway |
| | Apoptosis/tissue regeneration |
| ATF4 | Positive regulation of apoptotic process |
| JTB | Inhibits apoptosis induced by TGFB1 |
| SIVA1 | Viral entry into host cell apoptosis regulation |
| | Splicing |
| RBM8A | Regulation of alternative mRNA splicing via spliceosome |
| | Other |
| RPL4 | Viral process |
| RPL18A | Viral process |
| RPL22 | Viral process |
| RPLP1 | Viral process |
| RPS17 | Viral process |
| RPSAP58 | Ribosomal small subunit assembly |
| RPP25L | Ribonuclease P protein subunit p25-like protein |
| CUNH6orf226 | |

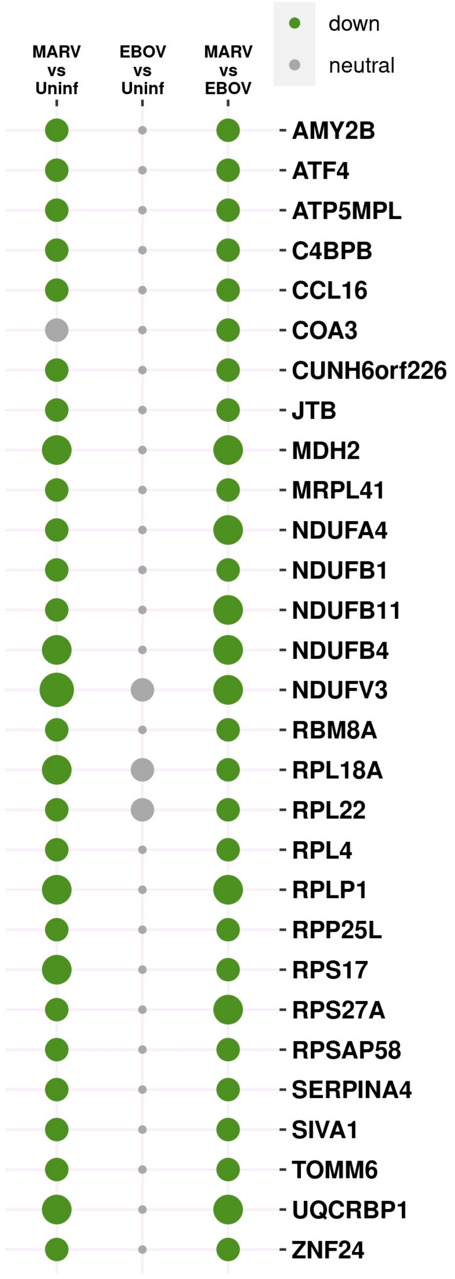


The balloon plot at the right compares responses of genes to MARV-, EBOV- infected and uninfected samples against each other. The radius of circle is proportional to $\log_2$ (ratio), red is for positive, green is for negative values and gray is used when absolute values of $\log_2$ (ratio) < 0.6.

**Table ST7. Divergent pathways and genes upregulated by EBOV but not MARV infection: vascular function, inflammation, mitochondria, lipid metabolism, tissue regeneration**

| Gene | Process |
|---|---|
| | Vascular |
| CYP11B2 | Regulation of blood volume by renal aldosterone Cytochrome P450 11B2 mitochondrial |
| TMEM133/ARHGAP42 | Inhibits RhoA activity to regulate vascular tone and control blood pressure |
| | Inflammation/stress |
| CYP11B1 | Cortisol production stress response Immune response Cytochrome P450 11B1 mitochondrial |
| | Mitochondrial |
| MRPS33 | Translation 28S ribosomal protein S33 |
| PET100 | Respiratory chain complex IV |
| NDUFA5 | Respiration electron transport NADH to Ubiquinone |
| | Lipid/fatty-acid |
| ADIRF | Lipid metabolism |
| | Tissue regeneration |
| H19 | lincRNA cell growth control |
| CENPW | Mitotic cell cycle |
| | Other |
| LINC00467 | lincRNA 467 |
| UNDEF312 | |

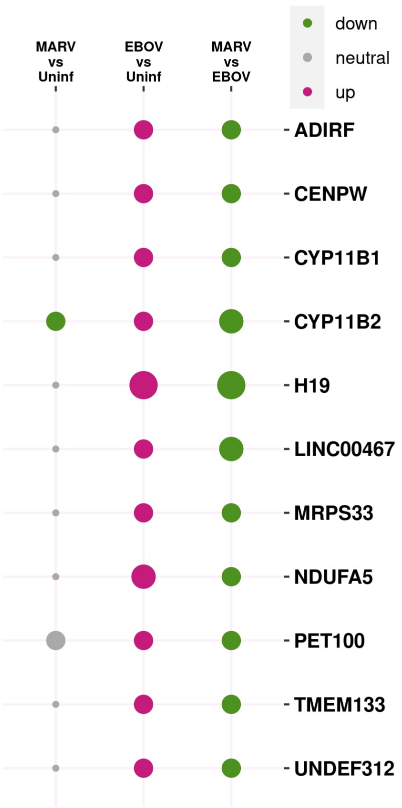


The balloon plot at the right compares responses of genes to MARV-, EBOV- infected and uninfected samples against each other. The radius of circle is proportional to $\log_2$ (ratio), red is for positive, green is for negative values and gray is used when absolute values of $\log_2$ (ratio) < 0.6.

**Table ST8. Divergent pathways and genes downregulated by EBOV but not MARV infection: innate immunity, coagulation and digestion**

| Gene | Process |
|---|---|
| | Innate immunity |
| **BST2** | Response to IFNG |
| | Vascular |
| **SERPINA13P** | Protease inhibitor clotting |
| | Digestion |
| **SLC51B** | Bile secretion |
| | Other |
| **UNDEF767** | |

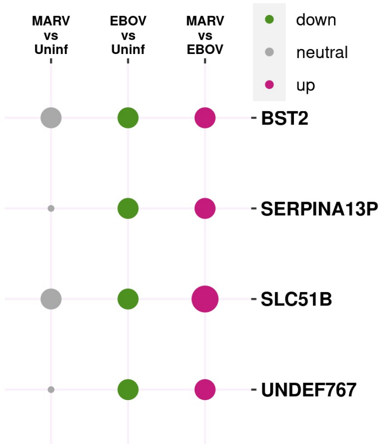


The balloon plot at the right compares responses of genes to MARV-, EBOV- infected and uninfected samples against each other. The radius of circle is proportional to $\log_2$ (ratio), red is for positive, green is for negative values and gray is used when absolute values of $\log_2$ (ratio) < 0.6.

#### Table ST9. Correlations between various samples based on expression levels of genes

**A. CB1 versus CB2 and cb3**. CB1 is dissimilar to CB2 and CB3

| This study | cb2_Uninf | cb3_Uninf | | ALL | cb2_Uninf | cb3_Uninf |
|---|---|---|---|---|---|---|
| cb1_Uninf | 0.77 | 0.74 | | cb1_Uninf | 0.73 | 0.77 |

**B. CB2 and cb3** are consistent with each other.

| This study | cb2_Uninf | cb3_Uninf | | ALL | cb2_Uninf | cb3_Uninf |
|---|---|---|---|---|---|---|
| cb2_Uninf | 1 | 0.94 | | cb2_Uninf | 1 | 0.91 |
| cb3_Uninf | 0.94 | 1 | | cb3_Uninf | 0.91 | 1 |

**C. CB versus MARV**. CB1 is similar to MARV while CB2 and cb3 are not.

| This study | ab06_mb | ab07_mb | ab08_mb | | ALL | ab06_mb | ab07_mb | ab08_mb |
|---|---|---|---|---|---|---|---|---|
| cb1_Uninf | 0.74 | 0.86 | 0.71 | | cb1_Uninf | 0.66 | 0.8 | 0.61 |
| cb2_Uninf | 0.26 | 0.46 | 0.24 | | cb2_Uninf | 0.31 | 0.5 | 0.27 |
| cb3_Uninf | 0.39 | 0.56 | 0.37 | | cb3_Uninf | 0.42 | 0.6 | 0.37 |

**D. CB versus EBOV.** CB2/3 are similar to EBOV while CB1 is not.

| This study | ab01_eb | ab02_eb | ab03_eb | | ALL | ab01_eb | ab02_eb | ab03_eb |
|---|---|---|---|---|---|---|---|---|
| cb1_Uninf | 0.81 | 0.86 | 0.69 | | cb1_Uninf | 0.82 | 0.85 | 0.58 |
| cb2_Uninf | 0.92 | 0.88 | 0.89 | | cb2_Uninf | 0.88 | 0.86 | 0.75 |
| cb3_Uninf | 0.99 | 0.96 | 0.98 | | cb3_Uninf | 0.96 | 0.93 | 0.79 |

**E. MARV versus MARV.** The MARV samples are consistent with each other.

| This study | ab06_mb | ab07_mb | ab08_mb | | ALL | ab06_mb | ab07_mb | ab08_mb |
|---|---|---|---|---|---|---|---|---|
| ab06_mb | 1 | 0.97 | 1 | | ab06_mb | 1 | 0.96 | 1 |
| ab07_mb | 0.97 | 1 | 0.96 | | ab07_mb | 0.96 | 1 | 0.94 |
| ab08_mb | 1 | 0.96 | 1 | | ab08_mb | 1 | 0.94 | 1 |

**F. EBOV versus MARV.** The correlations between MARV and EBOV samples are low

| This study | ab06_mb | ab07_mb | ab08_mb | | ALL | ab06_mb | ab07_mb | ab08_mb |
|---|---|---|---|---|---|---|---|---|
| ab01_eb | 0.47 | 0.64 | 0.45 | | ab01_eb | 0.5 | 0.68 | 0.45 |
| ab02_eb | 0.6 | 0.75 | 0.58 | | ab02_eb | 0.59 | 0.76 | 0.55 |
| ab03_eb | 0.38 | 0.54 | 0.36 | | ab03_eb | 0.35 | 0.49 | 0.33 |

**G. EBOV versus EBOV.** The EBOV samples are consistent with each other.

| This study | ab01_eb | ab02_eb | ab03_eb | | ALL | ab01_eb | ab02_eb | ab03_eb |
|---|---|---|---|---|---|---|---|---|
| ab01_eb | 1 | 0.99 | 0.96 | | ab01_eb | 1 | 0.96 | 0.78 |
| ab02_eb | 0.99 | 1 | 0.94 | | ab02_eb | 0.96 | 1 | 0.84 |
| ab03_eb | 0.96 | 0.94 | 1 | | ab03_eb | 0.78 | 0.84 | 1 |

In each sub-table, on the left are the correlations between samples considering only the genes used in this study and on the right are correlations calculated using all the genes detected by mRNAseq. These are consistent with the hypothesis that MARV infection of ERBs elicits a strong reaction, while EBOV infection is more muted and similar to the uninfected samples. cb1 shows hallmarks of inflammation, justifying its exclusion from the study (also seen in Fig 3). Uninf = uninfected, mb = MARV-infected, eb = EBOV-infected. Sample names start with cb or ab followed by numbers.
